# Structure of human endo-α-1,2-mannosidase (MANEA), an antiviral host-glycosylation target

**DOI:** 10.1101/2020.06.30.179523

**Authors:** Łukasz F. Sobala, Pearl Z Fernandes, Zalihe Hakki, Andrew J Thompson, Jonathon D Howe, Michelle Hill, Nicole Zitzmann, Scott Davies, Zania Stamataki, Terry D. Butters, Dominic S. Alonzi, Spencer J Williams, Gideon J Davies

## Abstract

Mammalian protein N*-*linked glycosylation is critical for glycoprotein folding, quality control, trafficking, recognition and function. N*-*linked glycans are synthesized from Glc_3_Man_9_GlcNAc_2_ precursors that are trimmed and modified in the endoplasmic reticulum (ER) and Golgi apparatus by glycoside hydrolases and glycosyltransferases. Endo-α-1,2-mannosidase (MANEA) is the sole *endo*-acting glycoside hydrolase involved in N*-*glycan trimming and unusually is located within the Golgi, where it allows ER escaped glycoproteins to bypass the classical N*-*glycosylation trimming pathway involving ER glucosidases I and II. There is considerable interest in the use of small molecules that disrupt N-linked glycosylation as therapeutic agents for diseases such as cancer and viral infection. Here we report the structure of the catalytic domain of human MANEA and complexes with substrate-derived inhibitors, which provide insight into dynamic loop movements that occur upon substrate binding. We reveal structural features of the human enzyme that explain its substrate preference and the mechanistic basis for catalysis. The structures inspired the development of new inhibitors that disrupted host protein N*-*glycan processing of viral glycans and reduced infectivity of bovine viral diarrhea and dengue viruses in cellular models. These results may contribute to efforts of developing broad-spectrum antiviral agents and bring about a more detailed view of the biology of mammalian glycosylation.

**SIGNIFICANCE STATEMENT:** The glycosylation of proteins is a major protein modification that occurs extensively in eukaryotes. Glycosidases in the secretory pathway that trim N-linked glycans play a key role in protein quality control and in the specific modifications leading to mature glycoproteins. Inhibition of glucosidases in the secretory pathway is a proven therapeutic strategy, and one with great promise in the treatment of viral disease. The enzyme endo-α-1,2-mannosidase, MANEA, provides an alternative processing pathway to evade glucosidase inhibitors. We report the 3D structure of human MANEA and complexes with enzyme inhibitors that we show act as antivirals for bovine viral diarrhea and human dengue viruses. The structure of MANEA will support inhibitor optimization and the development of more potent antivirals.

## Introduction

Asparagine (N) linked glycosylation is a protein modification that is widespread in eukaryotes and is essential for protein quality control and trafficking (1). N-Linked glycans contribute to protein structure and stability, receptor targeting, development and immune responses (2). N*-*Linked glycosylation occurs in the secretory pathway and is initiated in the lumen of the endoplasmic reticulum (ER) through co-translational transfer of the triantennary Glc_3_Man_9_GlcNAc_2_ glycan from a lipid linked precursor (Fig. 1; see also (2–5)). Formation of mature glycoproteins requires the removal of the glucose residues, typically through the action of α-glucosidases I and II, which are localized within the rough ER. However, in most cell lines and in human patients genetic disruption of α-glucosidase I or II, or inhibition of their activities do not prevent the formation of mature glycoproteins, because of the action of Golgi endo-α-1,2-mannosidase (MANEA (6)). The action of MANEA provides a glucosidase independent pathway for glycoprotein maturation termed the endomannosidase pathway (7).

**Figure 1.**
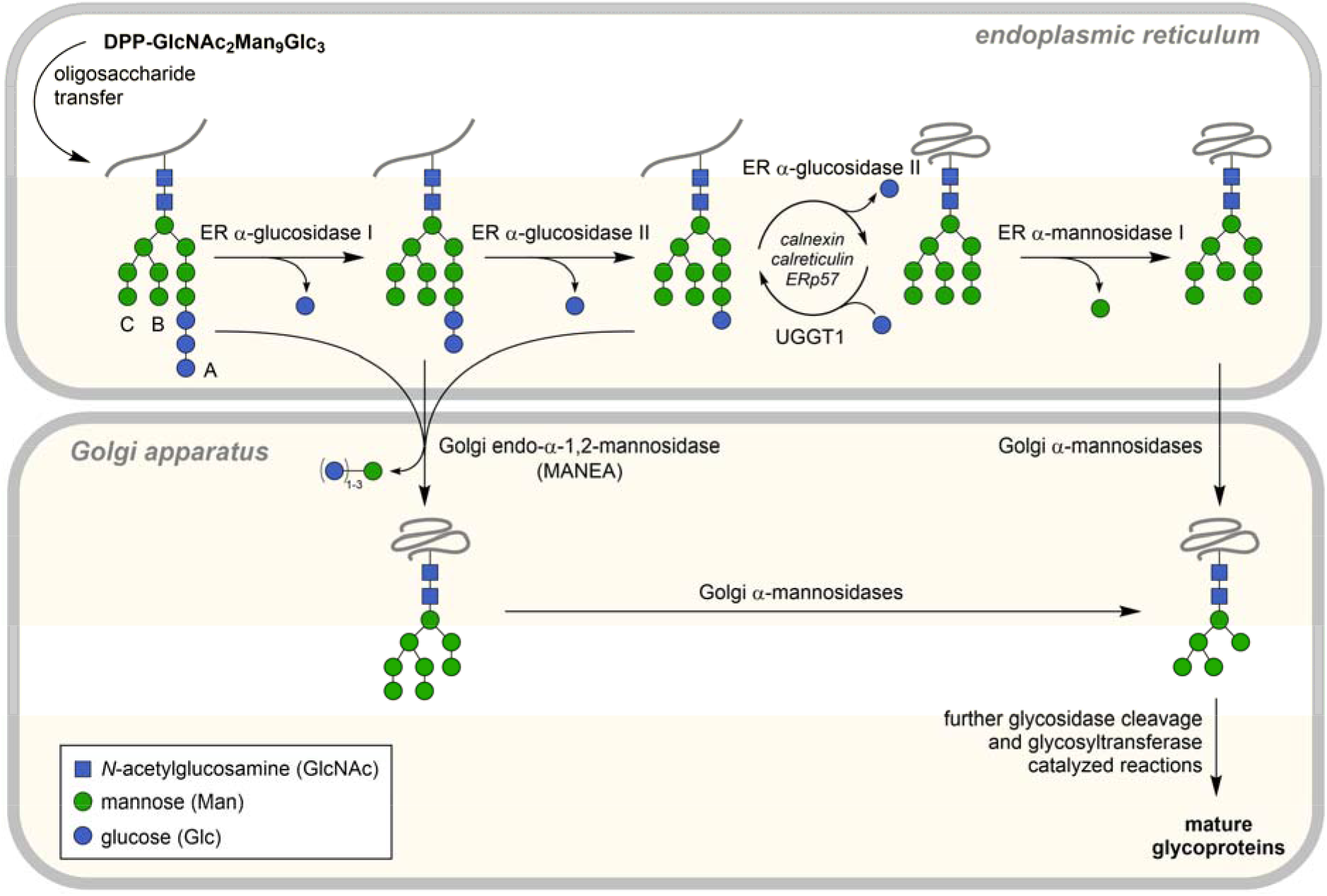
Simplified pathway for biosynthesis of N-linked glycans through the classical and endomannosidase pathways. *En bloc* transfer of the preformed Glc_3_Man_9_GlcNAc_2_ tetradecasaccharide (branches labelled A/B/C) from the dolichol-precursor occurs co-translationally to Asn residues within the consensus sequence Asn-Xxx-Ser/Thr. Trimming of glucose residues can be achieved through the classical pathway involving sequential action of α-glucosidases I and II. Alternatively, Golgi endo-α-1,2-mannosidase (MANEA) provides a glucosidase independent pathway for glycoprotein maturation, through cleaving the glucose-substituted mannose residues. ER mannosidase I resides in quality control vesicles (QCVs) (46). ER associated degradation of terminally misfolded glycoproteins is omitted.

**Figure 2.**
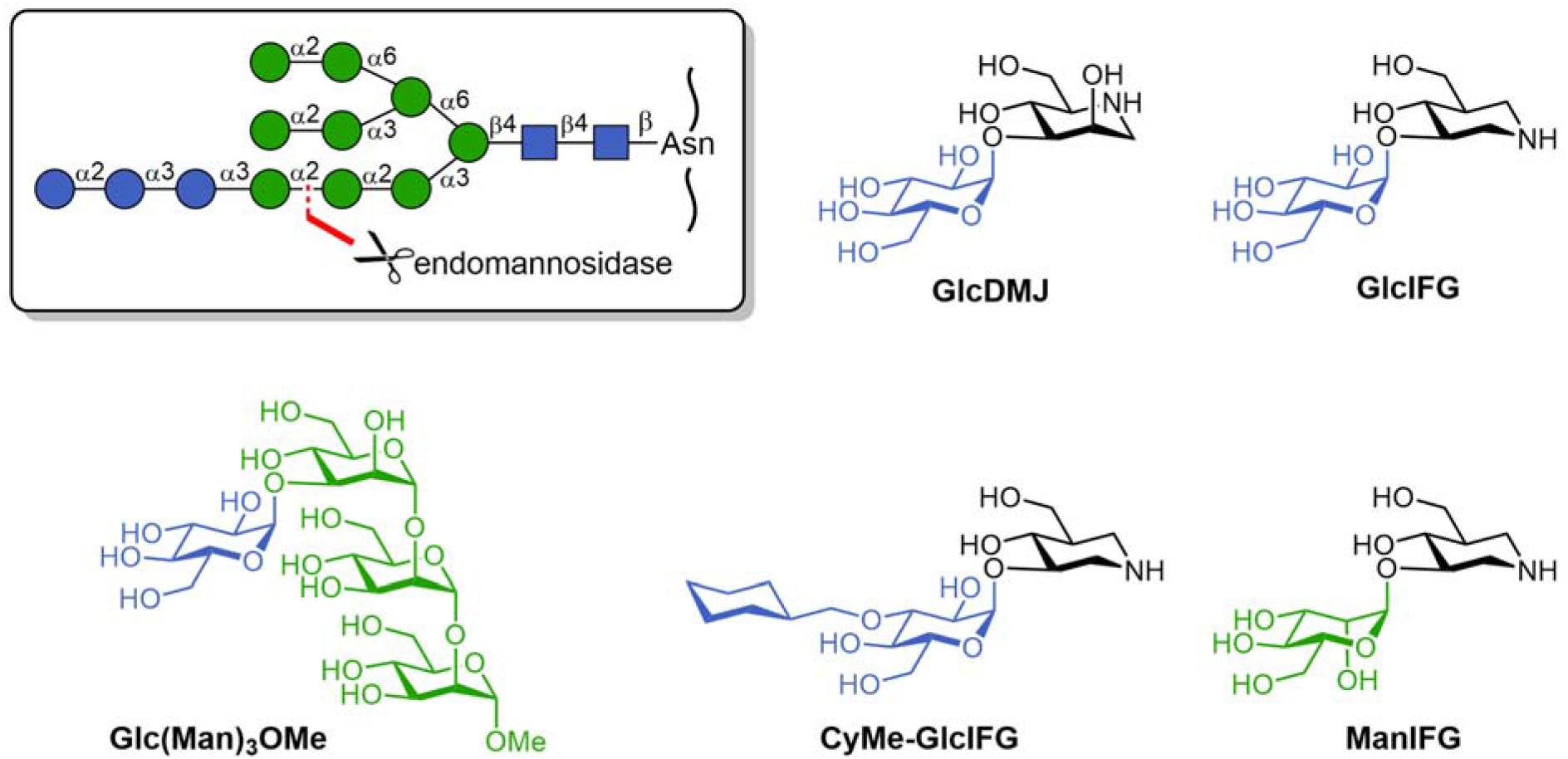
Substrates and inhibitors for MANEA inspired from the structure of the glucosylated N-glycan substrate. Inset: structure of Glc_3_Man_9_GlcNAc_2_ showing the cleavage site of MANEA.

MANEA is present chiefly in *cis*/*medial* Golgi (84%) and ER-Golgi intermediate compartment (ERGIC; 15%) (8) and allows trimming of glucosylated mannose residues from Glc_1-3_Man_9_GlcNAc_2_ structures and B and C branch mannose-trimmed variants (see Fig. 1 for definition of N-glycan branches). MANEA action on Glc_1-3_Man_9_GlcNAc_2_ results in cleavage of the glucosylated mannose in the A branch, releasing Glc_1-3_Man and Man_8_GlcNAc_2_; human MANEA acts faster on Glc_1_Man than on Glc_2,3_Man glycans (9). The endomannosidase pathway allows processing of ER-escaped, glucosylated high-mannose glycans on glycoproteins that fold independently of the calnexin/calreticulin (CNX/CRT) folding cycle, allowing them to rejoin the downstream N-glycan maturation pathway. In murine BW6147 cells under normal conditions, flux through the endomannosidase pathway accounts for around 15% of total flux through the secretory pathway (10); under conditions of glucosidase blockade through knockout or inhibition, MANEA can support as much as 50% of normal glycoprotein flux, depending on cellular expression levels (10). MANEA activity accounts for the accumulation of Glc3Man tetrasaccharide in the urine of a neonate suffering from CDG-IIb, a rare genetic disease caused by a deficiency of ER glucosidase I (MOGS) (11).

Most enveloped viruses (eg coronaviruses, retroviruses, ebolaviruses, hepatitis B virus (HBV), influenza viruses) contain glycoproteins within their envelope (12). Glycosylation is achieved by coopting host cell glycosylation machinery during replication. Substantial efforts have been deployed in the use of inhibitors of α-glucosidases I and II as antivirals (eg *N*-butyldeoxynojirimycin, 6-*O*-butanoylcastanospermine) for the treatment of HIV/AIDS (13), dengue (14, 15), and hepatitis B (16, 17) (reviewed in refs (18–20)). Inhibition of host glycosylation pathways, in particular ER glucosidases I and II and Golgi mannosidase I, interferes with the viral lifecycle by impairing protein folding and quality control, inducing the unfolded protein response, or mistrafficking of viral glycoproteins. These changes can lead to impairment of secretion (17, 21–23), fusion (24–26) or evasion of host immunity (27). In the case of HBV, when glucosidases are inhibited mature HBV viral glycoproteins are still produced through the alternative processing provided by the endomannosidase pathway (28). However, the contribution of the endomannosidase pathway to viral protein assembly under normal conditions remains poorly studied, and the antiviral activity of inhibitors targeting MANEA has not been evaluated.

The *Homo sapiens* endo-α-1,2-mannosidase gene, *MANEA*, is located on chromosome 6. The *MANEA* gene has one isoform whose protein product, MANEA, is categorized as a member of the glycoside hydrolase (GH) family 99 in the Carbohydrate-Active enZyme (CAZy) database (29), which also contains bacterial endo-α-1,2-mannanases from *Bacteroides xylanisolvens* (*Bx*GH99) and *Bacteroides thetaiotaomicron* (*Bt*GH99) that act on related structures within yeast mannans, and which share approx. 40% identity (30–32). *MANEA* encodes a protein 462 aa long that consists of a single pass, type II, membrane protein with a transmembrane helix followed by a stem region, followed by the catalytic domain (33). Here, we reveal the structure of the catalytic domain of human MANEA. Through complexes with substrate we reveal the architecture of the binding groove and identify key catalytic residues. We report structures with MANEA inhibitors that were designed based on the structure of the glucosylated high mannose N-glycan. Finally, we show that inhibitors of MANEA act as anti-viral agents against bovine viral diarrheal virus (BVDV) and dengue virus (DENV), confirming that MANEA is an anti-viral target.

## Results

### Expression and activity of Human MANEA GH99

In order to obtain soluble MANEA GH99 protein for structure-function studies to inform inhibitor design, attempts were made to express the gene. A truncated gene for MANEA, consisting of the catalytic domain beyond the stem domain, *i.e.* residues 98-462 (hereafter MANEA-Δ97), was synthesized in codon-optimized form. Expression trials in *E. coli* BL21(DE3) cells gave only insoluble protein, but we were inspired by a report (34) of soluble expression using cold-shock promotors (pCold-I vector) and co-expression of GroEL chaperones. Optimal yields of soluble soluble MANEA-Δ97 were obtained using Terrific Broth supplemented with glycerol and 20 mM MgCl_2_ affording approx. 2-3 mg L^−1^ (**Supplemental Figure 1A**, see Experimental procedures). The recombinant protein was purified using the encoded N-terminal His_6_-tag and was stable with a *T*_m_ of 50 ℃ (**Supplemental Figure 1B**).

Treatment of GlcMan_9_GlcNAc_2_ with recombinant MANEA-Δ97 released α-Glc-1,3-Man and Man_8_GlcNAc_2_ (**Supplemental Figure 1C**). Enzyme kinetics were measured using α-Glc-1,3-α-Man-1,2-α-Man-1,2-α-Man-OMe (GlcMan_3_OMe) as substrate using a coupled assay in which the product α-1,2-Man-ManOMe is hydrolysed by α-mannosidase and the resultant mannose quantified using a D-mannose/D-fructose/D-glucose detection kit (Megazyme, Inc.) (31). MANEA-Δ97 hydrolysed GlcMan_3_OMe with a *k*_cat_ of 27.7 ± 1.0 min^−1^ and *K*_M_ of 426 ± 33 μm (*k*_cat_/*K*_M_ of 65 mM^−1^ min^−1^) (**Supplemental Figure 1D**). The catalytic efficiency is similar to that displayed by *Bt*GH99 on the epimeric Man4OMe substrate (*K*_M_ = 2.6 mM, *k*_cat_ = 180 min^−1^, *k*_cat_/*K*_M_ = 69 mM^−1^ min^−1^) (31). The E404Q variant of MANEA-Δ97 was inactive on this substrate, consistent with the proposed mechanism (35).

### Development of MANEA inhibitors

GlcDMJ is a cell-permeable inhibitor of MANEA (36, 37) and is comprised of the well-known mannosidase iminosugar inhibitor deoxymannojirimycin (DMJ) modified with a glucosyl residue at the 3-position to enhance specificity and binding to MANEA by mimicking the substrate and benefitting from substrate-enzyme contacts in the -2 subsite. The related compound GlcIFG (38), which is also cell-permeable (39), was developed through a similar approach applied to the azasugar isofagomine (IFG). Similarly, ManIFG was synthesized to match the substrate stereochemistry of yeast mannan and is a superior inhibitor of bacterial endo-α-1,2-mannanases (31). As glucosylated mannans are substrates for α-glucosidase II, GlcDMJ and GlcIFG may potentially be degraded in cell-based systems (40), leading to loss of inhibition. Examination of the 3D X-ray structure of α-glucosidase II in complex with *N*-butyldeoxynojirimycin reveals binding of the sugar-shaped heterocycle involves a pocket-like active site that will not readily accommodate additional substituents (41). Conversely, it is known that MANEA can cleave Glc_1-3_Man substrates in which a substituent is tolerated at the 3-position of the glucosylated mannose. Therefore, we synthesized CyMe-GlcIFG, which bears a bulky cyclohexylmethyl substituent at the same position, in the expectation that it should be impervious to α-glucosidase II activity, yet still bind to MANEA.

Isothermal titration calorimetry revealed that GlcIFG binds to MANEA-Δ97 with *K*_d_ = 19.6 ± 5.6 nM and ManIFG with *K*_d_ = 170 ± 32 nM. Whilst we were encouraged by preliminary studies on a bacterial MANEA homolog that showed that CyMeGlc-IFG bound better (*K*_d_ = 284 nM) than GlcIFG alone (*K*_d_ = 625 nM), on the human enzyme CyMe-GlcIFG bound less tightly that GlcIFG with *K*_d_ = 929 ± 52 nM (**Supplemental Figure 1E**), consistent with structural features observed subsequently (see below). GlcDMJ, previously reported to have IC_50_ values of 1.7 to 5 μM (36, 40), was the weakest binding of the four compounds, with a *K*_d_ value of 1020 ± 36 nM. MANEA thus exhibits a preference for inhibitors that match the stereochemistry of the substrate −2/−1 residues (GlcIFG versus ManIFG), which is the reverse of the preference of bacterial endo-α-1,2-mannanases for the same inhibitors (i.e. for *Bt*GH99: GlcIFG *K*_d_ = 625 nM; ManIFG *K*_d_ = 140 nM (31)). Consistent with previous findings in bacterial GH99 enzymes, IFG is preferred to DMJ in the -1 subsite.

### Three-dimensional structure of human MANEA sheds light on eukaryotic enzyme specificity

Crystals of human MANEA-Δ97, **Figure 3A**, were obtained in several crystal forms (see Experimental). The initial crystal form, in space group *P*2_1_2_1_2, diffracted to around 2.25 Å resolution, and structure solution by molecular replacement (using the bacterial *Bx*GH99 as a search model) was successful leading to structures with *R*/*R*_free_ 18/22% (**Supplemental Table 1**). Unfortunately, this crystal form suffered from occlusion of the active centre by the His_6_-tag and a metal-ion assumed to be Ni^2+^. A second crystal form, *P*4_3_2_1_2 gave data of lower resolution (3 Å) but in this case we could build a loop (131–141) absent in the first crystal form; however, this crystal form could not be reliably reproduced. A third crystal form in space group *P*6_2_ was obtained with Anderson−Evans polyoxotungstate [TeW_6_O_24_]^6−^ (TEW), which has been used on occasion in protein crystallography to act as a linking agent between molecules in the crystal lattice (examples include PDB 4OUA, 4PHI, 4Z13, 6G3S, 6QSE and 6N90; reviewed in Ref (42)). The new crystal form diffracted up to 1.8 Å resolution, generating largely anisotropic datasets. The ligand binding site was occupied by HEPES, with residues 191-201 poised to accept –2/–1 subsite ligands and +1/+2 subsites open. The HEPES molecule was readily replaced by ligand soaking (the IC_50_ for HEPES shows weak binding, **Supplemental Figure S2**) and we obtained a series of complexes including a binary complex of MANEA-Δ97 with GlcIFG, a ternary complex with GlcIFG and α-1,2-mannobiose (structure and refinement statistics in **Supplemental Table 1**) and a complex of an inactive E404Q variant with a tetrasaccharide substrate.

**Figure 3.**
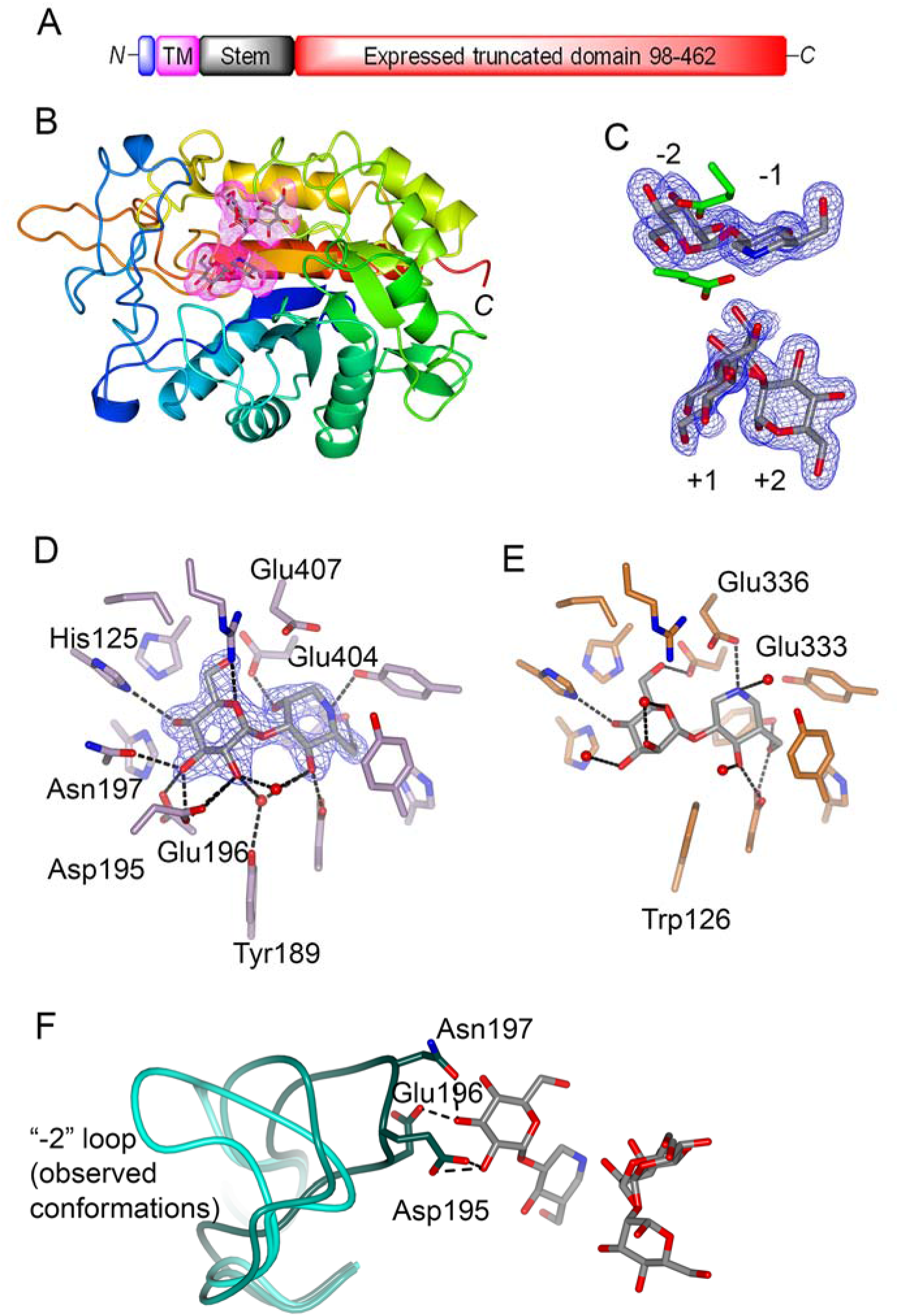
3D X-ray structure of *Homo sapiens* MANEA endomannosidase and its complexes. A. Domain structure of MANEA indicating the 98-462 domain that was expressed. B. 3D structure of MANEA, color ramped from N (blue) to C (red) terminus and with GlcIFG and α-1,2-mannobiose ligands shaded. C. Electron density for the ternary complex of MANEA with GlcIFG and α-1,2-mannobiose. 2m*Fo*-D*Fc* synthesis contoured at 0.4 e^−^/Å^3^. D. The –2 and –1 subsites of MANEA in complex with GlcDMJ, showing the interactions and with key residues labeled. 2m*Fo*-D*Fc* synthesis contoured at 0.6 e^−^/Å^3^. E. The comparable –2 and –1 subsites of the bacterial endomannanase *Bx*GH99 involved in yeast mannan degradation, in complex with ManIFG (PDB 4V27). F. Residues 189-203 containing the flexible loop as observed in: MANEA-E333Q structure (cyan), MANEA+Ni^2+^ structure (darker cyan) and MANEA with GlcIFG and α–1,2 mannobiose structure (darkest cyan; ligand: grey). Hydrogen bonds with the –2 sugar are shown as dashed lines.

The overall 3D structure of human MANEA is a single domain (*β/α*)_8_ barrel with a (partially, see below) open active center in which Glu404 and Glu407 (human numbering, as part of a conserved **E**WH**E**) motif are the catalytic residues in a neighboring group participation mechanism that proceeds through an epoxide intermediate (35), **Figure 3B,C**. The structure is similar (Cα RMSD over 0.9 Å over 333 matched residues) to structures reported (38) for *Bt*GH99 and *Bx*GH99, with which MANEA shares 40% sequence identity. The positions of the –2 to +2 ligands in the bacterial and human GH99 proteins are equivalent and the sugar interactions in the –2 to +2 subsites, all of which bind mannosides with identical linkages in both high-mannose N-glycans (MANEA) and yeast mannan (bacterial *endo*-α-1,2-mannanases (43)) are invariant.

A key difference between the substrates for the human and bacterial GH99 enzymes are the –2 sugar residues, which is glucose in high-mannose N-glycans, and mannose in yeast mannan. These differences are achieved by differences in recognition between the human and bacterial GH99 enzymes in the –2 subsite and its environs. In bacterial endo-α-1,2-mannanases Trp126 (numbering for *Bx*GH99 (38)) forms a hydrophobic interaction with the C2 that was believed to be responsible for the selectivity of bacterial enzymes for ManMan versus GlcMan substrates. In MANEA, the equivalent residue is Tyr189, which makes a water-mediated interaction with O2 of Glc in the –2 subsite, **Figure 3 D,E**. Notably, we observed a loop (residues 191-201, hereafter the “–2 loop)”, absent in the bacterial structures, that was flexible and observed in different position in the different MANEA crystal forms (**Figure 3F**). Two residues within the –2 loop are invariant across animal MANEAs: Asp195 and Gly198. Our structures reveal that Asp195 forms a hydrogen bond with the 3-OH of the –2 sugar (glucose) residue, and along with Asn197 H-bonding to O4, is a key determinant of binding of GlcMan structures. The second invariant residue, Gly198, enables the formation of a 195-198 turn thereby allowing residues 195, 196 and 197 to form hydrogen bonds with the –2 sugar (**Figure 3D,F)**.

MANEA isolated from human liver carcinoma cells processes triglucosylated N-glycans at a lower rate than monoglucosylated N-glycans (9). In the GlcIFG complex of MANEA, the –2 loop is closed over the active site, and in this conformation, triglucosylated N-glycans would be unable to bind to human MANEA. However, the other observed conformations show flexibility in the loop that could allow these extended structures to bind. Of note, bovine MANEA was observed not to process triglucosylated N-glycans (9). While both bovine and human MANEA possess the –2 loop, a Ser227 (human) to Lys change in the bovine enzyme was proposed contributed to the difference in specificity. Ser227 lies adjacent to, and points towards the –2 loop; the side chain of Lys226 (numbering as in bovine) may interact with the loop and reduce its mobility (**Figure 3F**). Interactions with the loop possibly explain why CyMe-GlcIFG binds nearly 50-fold more weakly than GlcIFG. Notably, bacterial GH99 endomannanases, which act on complex, extended yeast mannan substrates have open active sites and do not contain this loop (**Supplemental Figure 3**). To test this hypothesis, we determined the structure of *Bx*GH99 with CyMe-GlcIFG (*K*_d_ 339 nM; tighter binding than GlcIFG alone, 625 nM) and indeed the loop occludes the cyclohexyl binding and it is visible (but mobile) in the density (**Supplemental Figure 4**)

### Human MANEA as a host cell antiviral target

To explore the potential of the endomannosidase pathway as an antiviral target, we studied the effect of MANEA inhibitors on replication of BVDV, a pestivirus of the *Flaviviridae* family. Reinfection assays showed that increasing concentrations of GlcIFG, the tightest binding ligand of MANEA, resulted in a decrease in the number of infected cells (measured as focus-forming units (FFU)) **Figure 4A**. Previously, changes in the N-glycan structure of vesicular stomatitis virus G protein induced by the MANEA inhibitor GlcDMJ were identified by demonstrating a change in susceptibility to hydrolysis by endo-β-*N*-acetylglucosaminidase (endo H) (44). Therefore, we digested BVDV envelope glycoproteins E1/E2 with endo H. Increased sensitivity of E1/E2 proteins to endo H cleavage was observed both in the case of GlcIFG and the glucosidase I/II inhibitor NAP-DNJ, indicating an increased prevalence of high mannose N-glycans. The increased sensitivity for GlcIFG was greater than for NAP-DNJ, most likely reflecting the higher concentration of the former. Treatment with a combination of GlcIFG and NAP-DNJ gave an even higher sensitivity to endo H treatment, consistent with more effective cessation of N-glycan processing by blocking the glucosidase and endomannosidase pathways. We next examined BVDV replication in the presence of a combination of GlcIFG and NAP-DNJ (**Figure 4C**). These data showed that the combination of GlcIFG and NAP-DNJ gave a greater reduction than either agent alone suggesting an additive antiviral effect from inhibiting both the α-glucosidase I/II and endomannosidase pathways. The addition of NAP-DNJ not only inhibits calnexin mediated folding and quality control but requires viral glycans to undergo processing by MANEA, hence potentiating the antiviral effect.

**Figure 4.**
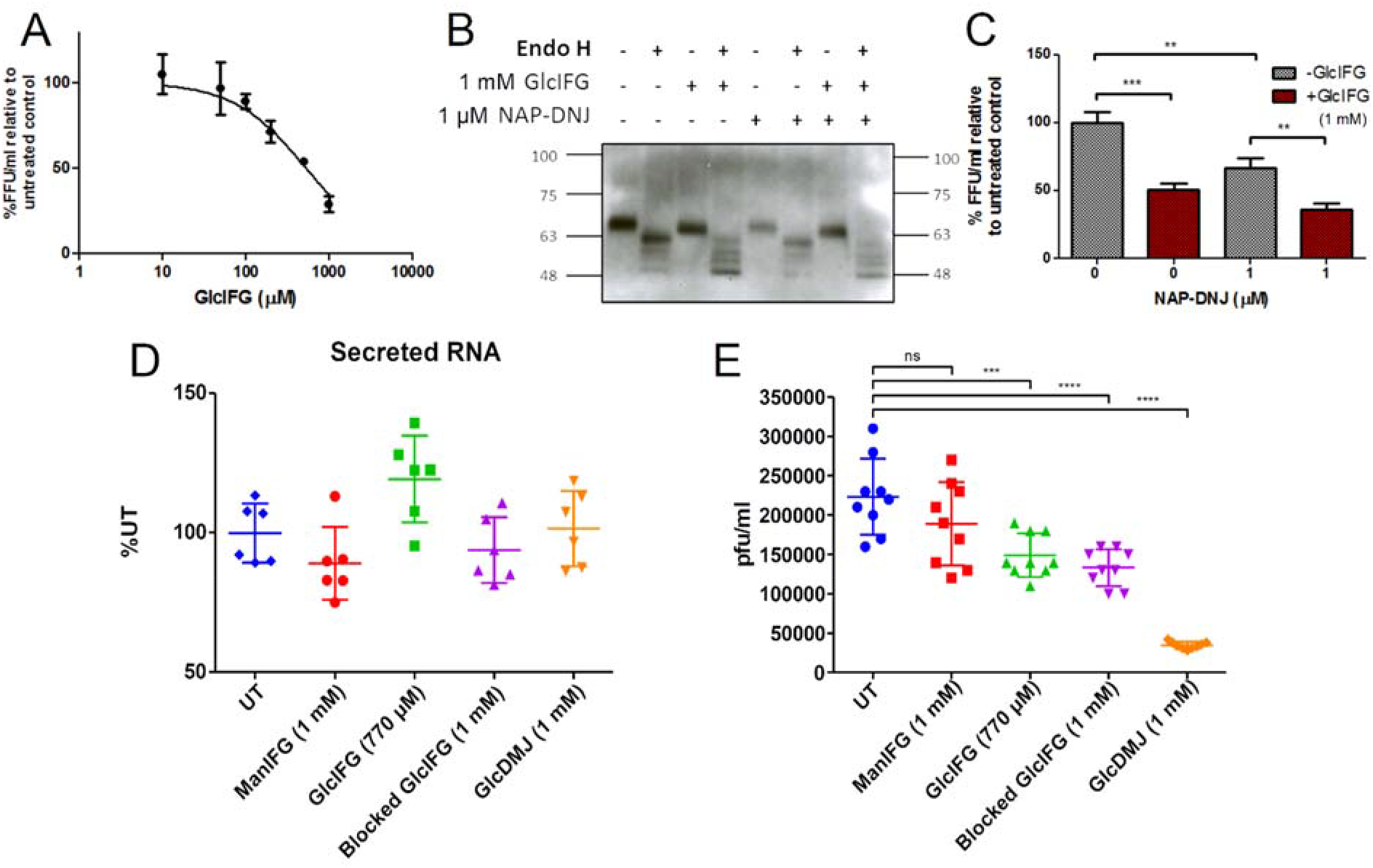
Anti-viral action of MANEA inhibitors. Results of (A,B,C) BVDV reinfection assays in MDBK cells and (D,E) DENV reinfection assays in Huh7.5 cells. (A) Percentage of FFU/ml relative to untreated cells at different concentrations of GlcIFG, at a multiplicity of infection of 1. (B) Effect of MANEA inhibition (GlcIFG) and ER glucosidase II inhibition (NAP-DNJ) on the susceptibility of glycans on the BVDV E1/E2 protein to cleavage by endoH. (C) The combined effects of NAP-DNJ and GlcIFG on BVDV infectivity, as measured by FFU/ml. Experiments performed in triplicate. (D) Secreted RNA levels in DENV infected Huh7.5 cells. (E) Reinfectivity plaque assay from DENV infected Huh7.5 cells. The horizontal bar in D and E indicates the mean.

To further define the antiviral potential of MANEA inhibitors, we extended these studies to DENV. The MANEA inhibitors GlcIFG, GlcDMJ, CyMe-GlcIFG and ManIFG were used to study levels of viral particle formation and infectivity. Viral particle formation, as assessed by secreted viral RNA, was unaffected by treatment with MANEA inhibitors (Figure 4D). However, treatment with GlcIFG, CyMe-GlcIFG and GlcDMJ caused a reduction in plaque number, showing that differences in glycosylation resulting from MANEA inhibition impair DENV infectivity. The greatest effect was seen for GlcDMJ, which reduced the number of plaque forming units by 6-fold; ManIFG had no effect. The antiviral effect observed is at a relatively high concentration but iminosugars are known to have difficulty in gaining access to the secretory pathway (21). This proof of principle demonstrates GlcDMJ is a potential broad-spectrum antiviral with changes in glycosylation reducing infectivity of the progeny. As it is the weakest-binding inhibitor, factors other than affinity, such as cell permeability, may be responsible for potency of antiviral effects.

### Summary

We report the 3D structure of Golgi endo-α-1,2-mannosidase, MANEA, a key eukaryotic N*-*glycosylation pathway glycosidase. This data provides a structural rationale to understand the change in specificity of this enzyme for mono-, di- and triglucosylated high mannose N-glycans. We also show the potential for MANEA inhibitors to alter N-glycan structures of viral envelope glycoproteins and reduce viral infectivity. MANEA processes Glc_1-3_Man_9_GlcNAc_2_ structures and provides a pathway for glycoprotein maturation that is independent of the classical α-glucosidase I/II dependent pathways. Treatment of HBV with miglustat (*N*-butyldeoxynojirimycin, an inhibitor of α-glucosidase II) impaired viral DNA secretion and led to aberrant N-glycans on M glycoprotein, but other viral glycoproteins displayed mature glycans that arose through the endomannosidase pathway (28). Conversely, GlcDMJ treatment of VSV led to changes in G protein glycosylation (44). These prior studies and the present work demonstrate that different viral glycoproteins have varying degrees of dependence on the α-glucosidase I/II and endomannosidase pathways for maturation, and that inhibition of MANEA alone can alter infectivity of two encapsulated viruses. The human MANEA 3D structure, alongside demonstrated anti-viral activity of disaccharide imino and aza sugar inhibitors, provides a foundation for future inhibitor and drug development work. Given the devastating consequences of global outbreaks of viral disease, the present work highlights the potential for MANEA as a new target for host-directed antiviral agents exploiting viral glycoprotein biosynthesis.

## ACKNOWLEDGEMENTS

We thank the European Research Council (ERC-2012-AdG-32294 “Glycopoise”), the Australian Research Council (DP120101396, FT130100103, DP180101957) and the Royal Society for the Ken Murray Research Professorship to GJD. We thank Diamond Light Source UK for access to beamlines I03, I04 and I24 (proposals mx1221, mx12587 and mx18598). We acknowledge Jon Agirre for the help with generating a dictionary file for TEW and Eleanor Dodson for assistance with processing of the highly anisotropic datasets. We thank Alexandra Males for collection of ITC data with Glc-DMJ and Mahima Sharma for help with supplemental figure production.

## METHODS

Full methods are provided as Supplemental Information.

### Expression, characterization and structure solution

Briefly, recombinant MANEA-Δ97 featuring an N-terminal His tag was expressed in *E. coli* from a pColdI system co-expressing the *groEL* and *groES* genes encoding the GroEL/ES chaperone system. Recombinant MANEA-Δ97 was purified by metal-ion affinity and cation-exchange chromatography. Extensive crystal screening identified five different crystal forms, with a condition including 1 mM TEW (tellurium-centered Anderson–Evans polyoxotungstate [TeW_6_O_24_]^6−^) allowing ligand binding studies without an occluded active site. Structure solution by molecular replacement, and refinement featured programs from the CCP4 suite (45). Structures, and observed data, have been deposited on the Protein Data Bank. MANEA activity was determined by mass spectrometry using GlcMan_9_GlcNAc_2_ as substrate with subsequent permethylation and analysis by MALDI-MS. Michael-Menten kinetics was performed using a coupled assay with GlcMan_3_OMe as substrate (31). Ligand binding thermodynamics were determined by isothermal titration calorimetry.

### Viral Infectivity Studies

MDBK cells were infected with BVDV at a multiplicity of infection (MOI) of 1, followed by incubation with different concentrations of GlcIFG (0-1 mM) and in combination with NAP-DNJ for 24 h. Cell culture medium was harvested and serial dilutions were made and used to infect naive MDBK cells. Cells were incubated for 24 h, then washed and fixed with 4% (v/v) paraformaldehyde in PBS for 30 min. Following washing and blocking, cells were permeabilised and incubated with MAb103/105 (1:500 dilution in 5% milk-PBS; Animal Health Veterinary Laboratories Agency, Weybridge, UK) for 1 h. Incubation with anti-mouse FITC-conjugated secondary antibody (1:500 dilution in 5% milk-PBS; Sigma) and staining with 4′,6-diamidino-2-phenylindole allowed counting of fluorescent foci and calculation of virus titers (fluorescent focus units (FFU)/ml). The samples obtained after treatment for 24 h were treated with Endo H and separated by SDS–PAGE under non-reducing conditions. The membrane produced following transfer was probed with MAb214 primary antibody and then visualized with anti-mouse horseradish peroxidase secondary antibody (1:1000; Dako) for 1 h.

Huh7.5 cells were infected for 2 h with 50 μL of DENV2 strain 16681 at an MOI of 0.1. The inoculum was removed and replaced with 200 μL of growth media with drug at concentrations as indicated in triplicate, and the infected cells were incubated with drug for 2 d. The supernatant was harvested. DENV RNA in cell culture supernatants was isolated according to the manufacturer’s protocol for Direct-zol RNA MiniPrep Kit (Zymo Research) and assayed by qRT-PCR. Samples were read in technical duplicate and compared to a standard curve generated from high-titer viral RNA isolated from C6/36-grown DENV2. 95% confidence intervals were determined based on biological and technical variation and graphed using Prism 6 (GraphPad Software, Inc). The infectious DENV titres in supernatants collected were evaluated by plaque assay.

