## Supplementary information for "Structure of human endo-α-1,2-mannosidase (MANEA), an antiviral host-glycosylation target"

**This PDF file includes:**

Figures S1 to S4

Tables S1 to S3

Supplemental Experimental Procedures

Supplemental References

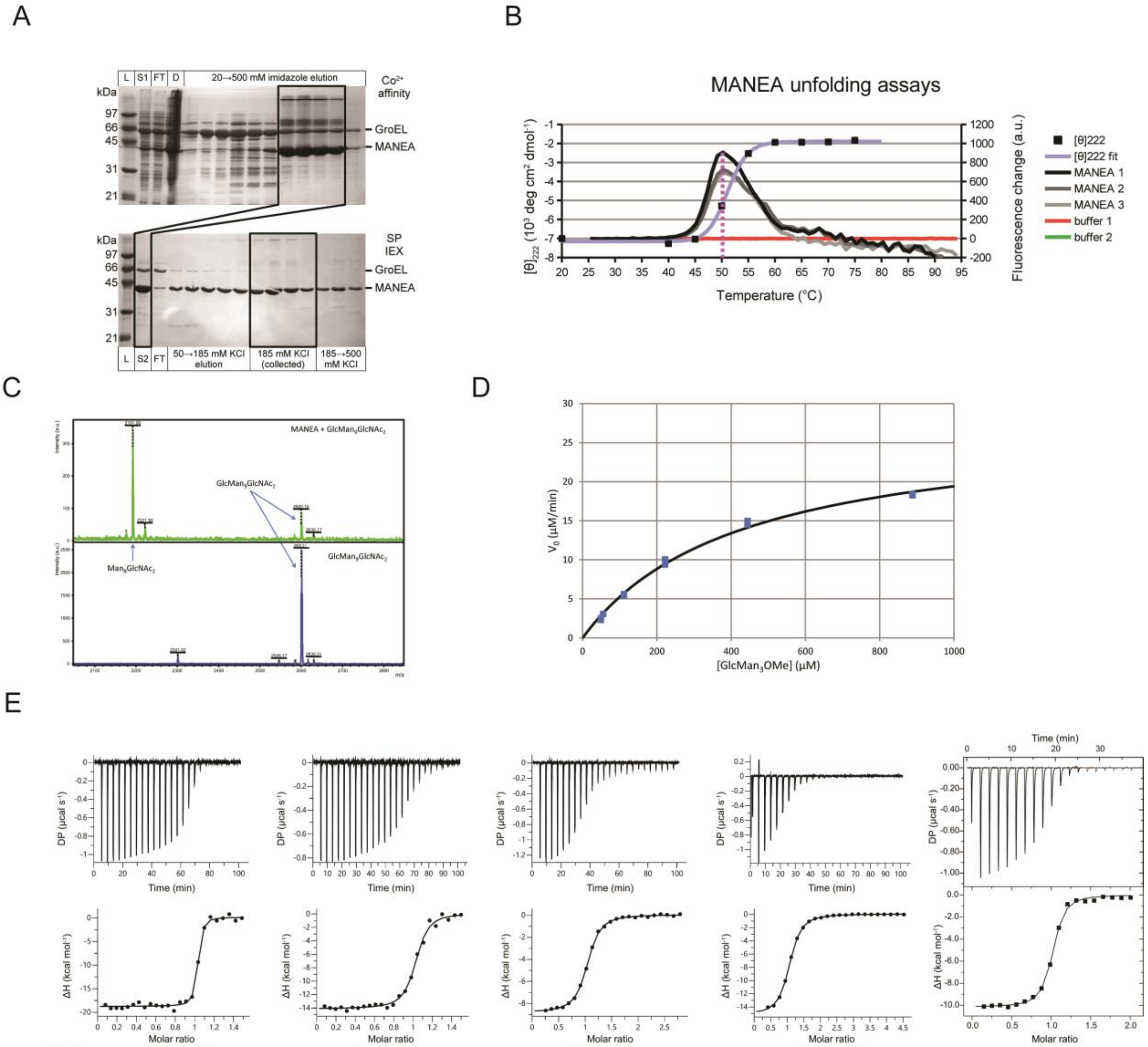

| Protein | MANEA-Δ97 | MANEA-Δ97 | MANEA-Δ97 | MANEA-Δ97 | *Bx*GH99 |
|---|---|---|---|---|---|
| Inhibitor | GlcIFG | ManIFG | CyMe-GlcIFG | GlcDMJ | CyMe-GlcIFG |
| N | 1 (fixed) | 1 (fixed) | 0.998 ± 0.003 | 1 ± 0.003 | 0.95 ± 0.004 |
| **$K_d$ (nM)** | **19.6 ± 5.6** | **170 ± 32** | **929 ± 52** | **1020 ± 36** | **339 ± 103** |
| $\Delta H$ (kcal mol$^{-1}$) | -18.9 ± 0.3 | -14.2 ± 0.4 | -8.85 ± 0.06 | -15.3 ± 0.09 | -10.3 ± 0.29 |
| $-T\Delta S$ (kcal mol$^{-1}$) | 8.35 | 4.96 | 0.624 | 7.09 | 5.07 |

**Supplemental Figure 1.**
**MANEA-Δ97 purification and assays.** (A) 12% SDS-PAGE of the purification procedure, top: 5 ml cobalt affinity column, bottom: 5 ml SP cation exchange column. Fractions highlighted in the top gel are representative of the pooled sample used for cation exchange (S2). Abbreviations used – L: protein ladder, S1/S2: samples loaded onto the columns, FT: column flow-through, D: lysed cell debris. (B) MANEA-Δ97 denaturation assays by circular dichroism (monitoring ellipticity at 222 nm – [θ]222) and thermal shift assay (SYPRO Orange fluorescence at 570 nm). (C) MANEA-Δ97 activity on a purified $GlcMan_9GlcNAc_2$ oligosaccharide. Top: $GlcMan_9GlcNAc_2$ and enzyme, bottom: $GlcMan_9GlcNAc_2$ alone. (D) Michaelis-Menten curve for MANEA-Δ97 and $GlcMan_3OMe$. (E) Isothermal titration calorimetry results for MANEA-Δ97 with GlcIFG, ManIFG & CyMe-GlcIFG, GlcDMJ and *Bx*GH99 with CyMe-GlcIFG.

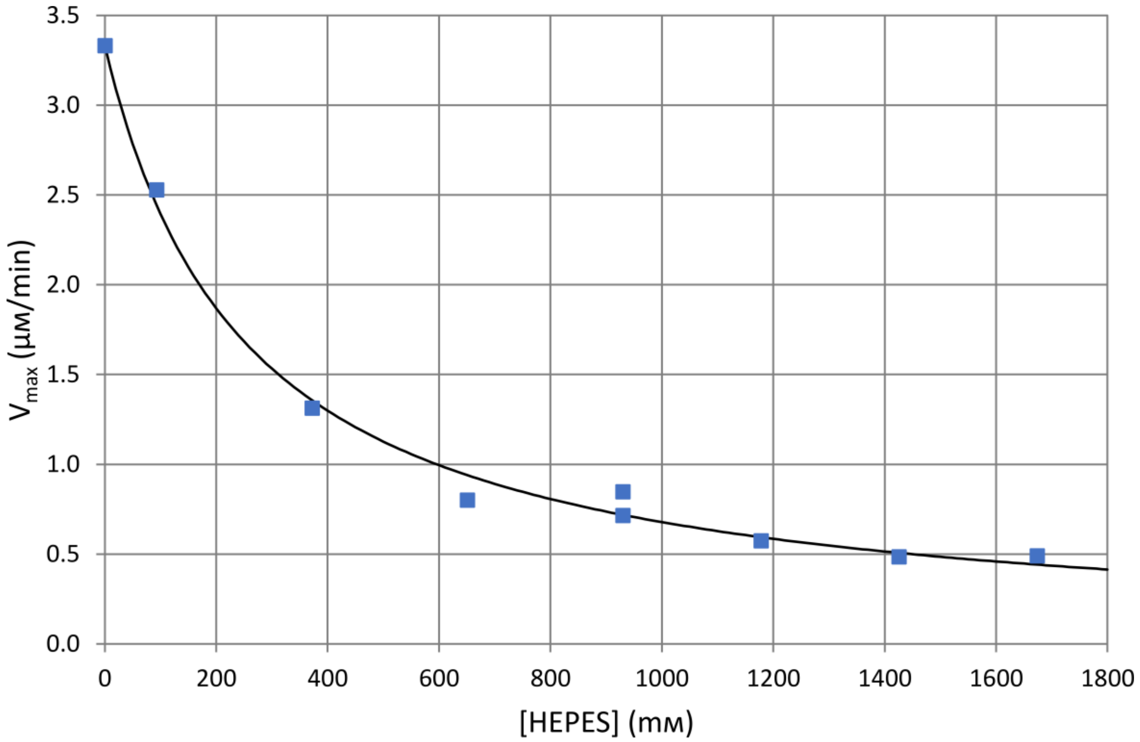

**Supplemental Figure 2.**

**HEPES inhibition.** $IC_{50}$ curve for HEPES inhibition of MANEA-Δ97. The concentration of the enzyme was 1 µM and the substrate was $GlcMan_3OMe$ (100 µM). The $IC_{50}$ value was 255 ± 12 mM ($R^2$=0.994). Due to a limited amount of available substrate, the experiments were done in single runs (except one HEPES concentration, as visible on the graph).

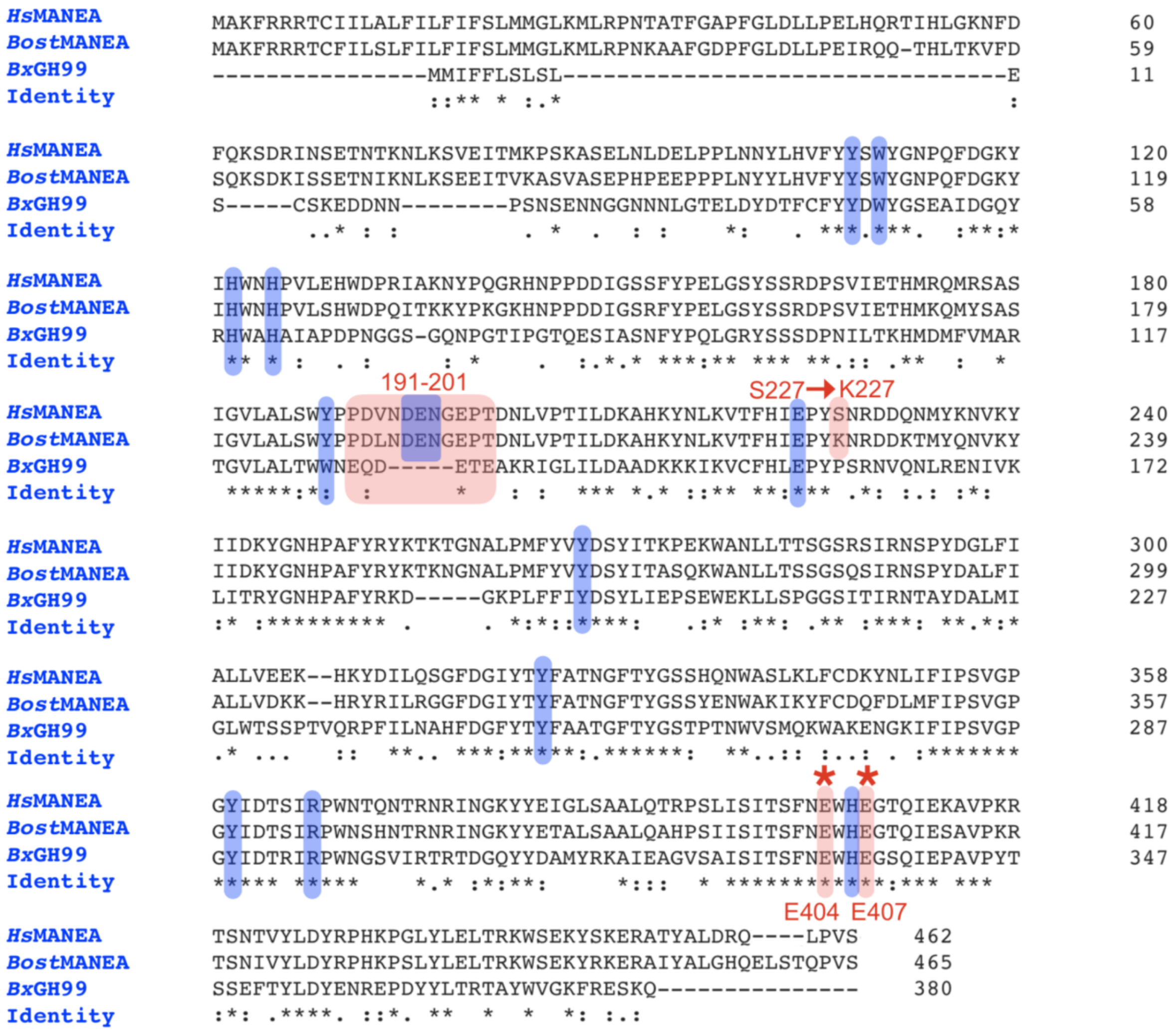

**Supplemental Figure 3.**

Multiple sequence alignment of GH99 MANEA enzymes from *Homo sapiens* (*Hs*), *Bos taurus* (Bovine, *Bost*) and *Bacteroides xylanisolvens (Bx)*. The -2 loop region 191-201, the catalytic dyad Glu404 and Glu407 in *Homo sapiens* and the Ser227→Lys227 in *Bos taurus* mutation are all highlighted in red (human numbering). Residues that make contacts with the –2 to +2 sugars are highlighted in blue. Sequence alignment was carried out using CLUSTAL Omega (1.2.4) MSA tool with manual correction.

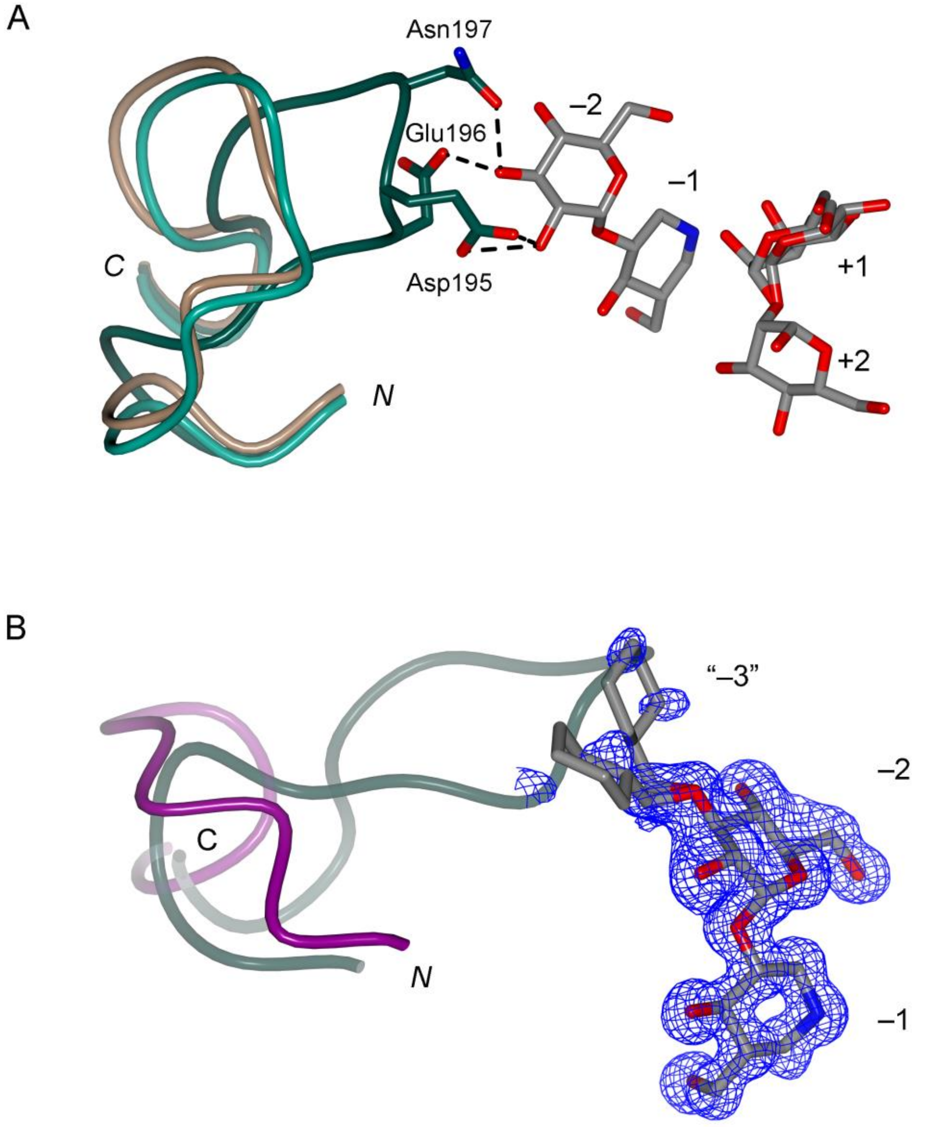

**Supplemental Figure 4.**

**Further details of the MANEA –2 loop.** (**A**) Comparison between the –2 loop in the wild-type MANEA-Δ97 ternary complex with GlcIFG and α-1,2-mannobiose (dark cyan), the $Ni^{2+}$ crystal form (cyan) and the bistris form (pale brown). In case of the bistris form the loop was more mobile and was modelled *de novo* at 50% occupancy resulting in a similar conformation to the $Ni^{2+}$ form. (**B**) Comparison of the residues aligned to those of the –2 loop in the structure of *Bx*GH99 with CyMe-GlcIFG (protein: purple, residues 127-135, inhibitor: grey, 2m*Fo*-D*Fc* synthesis contoured at 0.3 $e^{-}/Å^{3}$) and the relative position of the loop in structurally aligned MANEA-Δ97 ternary complex with GlcIFG and α-1,2-mannobiose (dark cyan with transparency, residues 189-203). Figure made in ccp4mg.

**A.**

<u>CATATG</u>CCGCTGAACAATTACCTGCATGTGTTTTATTACTCCTGGTATGGCAACCCGCAGTT
CGATGGTAAATACATTCATTGGAATCACCCGGTTCTGGAACATTGGGACCCGCGTATCGCTA
AAAACTATCCGCAAGGCCGCCACAATCCGCCGGATGACATTGGCAGCTCTTTTTACCCGGAA
CTGGGTAGCTATAGTTCCCGCGATCCGTCTGTTATTGAAACGCACATGCGTCAGATGCGCAG
CGCATCTATCGGCGTCCTGGCTCTGAGCTGGTATCCGCCGGATGTGAACGACGAAAATGGTG
AACCGACGGATAACCTGGTTCCGACCATTCTGGACAAAGCGCATAAATATAACCTGAAAGTC
ACCTTCCACATCGAACCGTACTCAAACCGTGATGACCAAAACATGTACAAAAACGTCAAATA
CATCATCGATAAATACGGCAACCATCCGGCGTTTTATCGCTACAAAACCAAAACGGGTAATG
CCCTGCCGATGTTCTATGTGTACGACAGCTATATCACCAAACCGGAAAAATGGGCGAACCTG
CTGACCACGAGTGGCTCCCGTTCAATTCGCAATTCACCGTATGATGGTCTGTTTATCGCCCT
GCTGGTTGAAGAAAAACATAAATACGATATCCTGCAGTCGGGCTTCGACGGTATCTATACCT
ACTTTGCAACGAACGGCTTCACCTATGGTTCATCGCACCAAAATTGGGCTAGTCTGAAACTG
TTTTGCGATAAATACAACCTGATCTTCATCCCGAGTGTCGGCCCGGGTTACATTGACACCTC
CATCCGTCCGTGGAACACCCAGAATACGCGTAACCGCATTAATGGCAAATATTACGAAATTG
GTCTGTCTGCCGCACTGCAGACCCGTCCGTCTCTGATTAGTATCACCTCCTTTAACGAATGG
CATGAAGGCACGCAAATTGAAAAAGCCGTGCCGAAACGTACCAGCAATACGGTTTATCTGGA
TTACCGCCCGCACAAACCGGGTCTGTATCTGGAACTGACCCGTAAATGGTCTGAAAAATACT
CAAAAGAACGTGCCACCTATGCTCTGGATCGTCAACTGCCGGTCTCCTGA<u>CTCGAG</u>

**B.**

**MNHKVHHHHHHIEGRH**MPLNNYLHVFYYSWYGNPQFDGKYIHWNHPVLEHWDPRIAKNYPQG
RHNPPDDIGSSFYPELGSYSSRDPSVIETHMRQMRSASIGVLALSWYPPDVNDENGEPTDNL
VPTILDKAHKYNLKVTFHIEPYSNRDDQNMYKNVKYIIDKYGNHPAFYRYKTKTGNALPMFY
VYDSYITKPEKWANLLTTSGSRSIRNSPYDGLFIALLVEEKHKYDILQSGFDGIYTYFATNG
FTYGSSHQNWASLKLFCDKYNLIFIPSVGPGYIDTSIRPWNTQNTRNRINGKYYEIGLSAAL
QTRPSLISITSFNEWHEGTQIEKAVPKRTSNTVYLDYRPHKPGLYLELTRKWSEKYSKERAT
YALDRQLPVS*

**Supplemental Figure 5.**

**DNA and protein sequences.** (**A**) DNA sequence of the *MANEA-Δ97* gene as cloned into pCold-I vector. Restriction endonuclease sites (*NdeI* and *XhoI*) are underlined. (**B**) Full amino acid sequence of MANEA-Δ97 protein. Expression tag residues are bolded.

**Supplemental Table S1.** Data collection and model refinement statistics of the MANEA-Δ97 structures discussed in the article. *Values in parentheses represent the highest resolution shell.

| | WT with $Ni^{2+}$ | WT with HEPES | WT with HEPES and TEW | WT with GlcIFG and TEW | WT with GlcIFG, α-1,2-mannobiose and TEW | WT with bis-tris | E404Q with αMan-1,2-ManOMe |
|---|---|---|---|---|---|---|---|
| Crystal form | 1 | 2 | 3 | 3 | 3 | 4 | 5 |
| **Data collection** | | | | | | | |
| Diamond beamline | I03 | I04 | I03 | I03 | I03 | I24 | I03 |
| Space group | $P2_12_12$ | $P4_32_12$ | $P6_2$ | $P6_2$ | $P6_2$ | $P2_12_12_1$ | $P2_1$ |
| Cell dimensions | | | | | | | |
| *a*, *b*, *c* (Å) | 38.5, 86.5, 135.9 | 144.2, 144.2, 139.7 | 129.0, 129.0, 48.8 | 127.9, 127.9, 48.8 | 127.9, 127.9, 48.4 | 42.8, 107.0, 43.9 | 42.7, 81.7, 53.0 |
| α, β, γ (°) | 90, 90, 90 | 90, 90, 90 | 90, 90, 120 | 90, 90, 120 | 90, 90, 120 | 90, 90, 90 | 90, 92.93, 90 |
| Resolution (Å)* | 135.92–2.25 (2.33–2.25) | 102.00–3.00 (3.16–3.00) | 64.48–2.00 (2.05–2.00) | 31.97–1.90 (1.94–1.90) | 110.76–1.80 (1.84–1.80) | 38.41–1.20 (1.22–1.20) | 52.92–1.10 (1.12–1.10) |
| $R_{merge}$ | 0.200 (1.872) | 0.596 (1.175) | 0.126 (0.739) | 0.203 (3.430) | 0.148 (0.939) | 0.049 (0.695) | 0.054 (0.223) |
| $R_{pim}$ | 0.083 (1.002) | 0.135 (0.265) | 0.078 (0.450) | 0.050 (0.843) | 0.038 (0.237) | 0.028 (0.485) | 0.029 (0.150) |
| *CC(1/2)* | 0.994 (0.345) | 0.974 (0.253) | 0.992 (0.411) | 0.998 (0.546) | 0.999 (0.782) | 0.999 (0.806) | 0.998 (0.944) |
| $\langle I / \sigma I \rangle$ | 7.0 (0.9) | 5.0 (2.9) | 6.2 (1.5) | 10.5 (1.0) | 13.4 (2.9) | 10.6 (1.5) | 11.9 (3.3) |
| Completeness (%) | 98.1 (84.5) | 100 (99.8) | 99.6 (98.3) | 100 (99.8) | 100 (100) | 99.4 (94.7) | 97.6 (71.3) |
| Redundancy | 11.4 (6.9) | 20.3 (20.5) | 3.5 (3.5) | 17.6 (17.4) | 16.5 (16.6) | 3.8 (2.8) | 3.7 (2.6) |
| **Refinement** | | | | | | | |
| Resolution (Å) | 73.10–2.25 | 102.00–3.00 | 64.48–2.00 | 31.97–1.90 | 110.76–1.80 | 38.44–1.20 | 52.98–1.10 |
| No. reflections all / free | 21817 / 1106 | 30099 / 1429 | 31481 / 1554 | 36184 / 1779 | 42104 / 2094 | 114455 / 5555 | 143263 / 7225 |
| $R_{work}$ / $R_{free}$ | 0.18 / 0.22 | 0.21 / 0.27 | 0.20 / 0.23 | 0.19 / 0.22 | 0.18 / 0.22 | 0.14 / 0.18 | 0.11 / 0.13 |
| No. atoms | | | | | | | |
| Protein | 2906 | 8853 | 2998 | 2961 | 2974 | 2949 | 3220 |
| Ligand/ion | 1 | 45 | 139 / 1 | 129 / 1 | 163 / 1 | 14 | 29 |
| Water | 185 | 140 | 165 | 113 | 137 | 456 | 491 |
| B-factors ($Å^2$) | | | | | | | |
| Protein | 51 | 49 | 34 | 32 | 26 | 25 | 15 |
| Ligand/ion | 88 | 52 | 76 / 47 | 66 / 40 | 47 / 29 | 38 | 19 |
| Water | 51 | 36 | 36 | 32 | 30 | 24 | 30 |
| RMS. deviations | | | | | | | |
| Bond lengths (Å) | 0.015 | 0.012 | 0.015 | 0.017 | 0.017 | 0.013 | 0.018 |
| Bond angles (°) | 1.9 | 1.7 | 2.0 | 2.1 | 2.2 | 1.8 | 2.1 |
| **PDB ID** | **6ZDC** | **6ZDF** | **6ZDK** | **6ZDL** | **6ZFA** | **6ZFQ** | **6ZFN** |

**Supplemental Table S2**. Data collection and model refinement statistics of the *Bx*GH99 structure with CyMe-GlcIFG. *Values in parentheses represent the highest resolution shell.

| | *Bx*GH99 with CyMe-GlcIFG |
|---|---|
| **Data collection** | |
| Diamond beamline | I03 |
| Space group | *I*4 |
| Cell dimensions | |
| *a*, *b*, *c* (Å) | 108.0, 108.0, 67.3 |
| α, β, γ (°) | 90, 90, 90 |
| Resolution (Å)* | 76.35–1.09 (1.11–1.09) |
| $R_{\text{merge}}$ | 0.059 (1.120) |
| $R_{\text{pim}}$ | 0.027 (0.715) |
| *CC(1/2)* | 0.997 (0.358) |
| $<I / \sigma I>$ | 12.5 (0.9) |
| Completeness (%) | 95.9 (66.4) |
| Redundancy | 5.1 (2.8) |
| **Refinement** | |
| Resolution (Å) | 76.35–1.09 |
| No. reflections all / free | 153946 / 7638 |
| $R_{\text{work}}$ / $R_{\text{free}}$ | 0.11 / 0.13 |
| No. atoms | |
| Protein | 3175 |
| Ligand/ion | 55 |
| Water | 468 |
| B-factors (Å$^2$) | |
| Protein | 17 |
| Ligand/ion | 23 |
| Water | 34 |
| RMS. deviations | |
| Bond lengths (Å) | 0.015 |
| Bond angles (°) | 1.9 |
| **PDB ID** | **6ZJ6** |

**Supplemental Table S3.** Data collection and model refinement statistics of the MANEA-Δ97 structures discussed in the article which were solved using anisotropically truncated datasets. *Values in parentheses represent the highest resolution shell.

| | WT with GlcDMJ and TEW | E404Q with $GlcMan_3OMe$ and TEW |
|---|---|---|
| Crystal form | 3 | 3 |
| **Data collection** | | |
| Diamond beamline | I04 | I04 |
| Space group | $P6_2$ | $P6_2$ |
| Cell dimensions | | |
| *a*, *b*, *c* (Å) | 128.5, 128.5, 48.3 | 129.4, 129.4, 50.2 |
| α, β, γ (°) | 90, 90, 120 | 90, 90, 120 |
| Diffraction limits | | |
| *a*, *b*, *c* (Å) | 2.27, 2.27, 4.02 | 1.96, 1.96, 3.43 |
| Resolution (Å)* | 111.33–2.27 (2.54–2.27) | 112.07–1.96 (2.15–1.96) |
| $R_{merge}$ | 0.276 (1.520) | 0.093 (1.448) |
| $R_{pim}$ | 0.103 (0.547) | 0.029 (0.463) |
| *CC(1/2)* | 0.989 (0.569) | 0.999 (0.658) |
| $\langle I / \sigma I \rangle$ | 5.3 (1.5) | 15.7 (1.6) |
| Completeness (%) | | |
| spherical | 51.9 (15.4) | 50.1 (11.9) |
| ellipsoidal | 88.8 (61.8) | 85.5 (50.6) |
| Redundancy | 8.0 (8.6) | 11.7 (10.5) |
| **Refinement** | | |
| Resolution (Å) | 111.33–2.27 | 64.71–1.96 |
| No. reflections all / free | 11089 / 545 | 17488 / 844 |
| $R_{work}$ / $R_{free}$ | 0.19 / 0.26 | 0.18 / 0.25 |
| No. atoms | | |
| Protein | 2993 | 3031 |
| Ligand/ion | 160 / 1 | 108 / 1 |
| Water | 94 | 133 |
| B-factors ($Å^2$) | | |
| Protein | 31 | 43 |
| Ligand/ion | 74 / 36 | 67 / 54 |
| Water | 22 | 40 |
| RMS. deviations | | |
| Bond lengths (Å) | 0.006 | 0.009 |
| Bond angles (°) | 1.5 | 1.8 |
| **PDB ID** | **6ZJ5** | **6ZJ1** |

### Molecular and Structural Biology

#### Cloning and mutagenesis of the human MANEA

A DNA sequence of the open reading frame (ORF) of human *MANEA* gene was optimized *in silico* for expression in *E. coli* and ordered from GenScript, Inc (**Supplementary Figure 4**). The sequence encoded residues 34-462 of the protein and was subcloned into pET28a (+) vector using *NdeI* and *XhoI* restriction sites. Subsequently, the sequence was truncated using PCR (Q5 High-Fidelity DNA Polymerase, NEB) in order to encode residues 98-462 and cloned into pColdI (1) vector (TaKaRa Bio, Inc.), which encodes an N-terminal $His_6$-tag, using *NdeI* and *XhoI* restriction sites,. The sequence of the forward primer used for the truncation was: GATCAG<u>CATATG</u>CCGCTGAACAATTACCTGCA and of the reverse primer: GATCAG<u>CTCGAG</u>TCAGGAGACCGG (the restriction endonuclease sites are underlined). The ORF encoding residues 98-462 of *MANEA* was fully Sanger sequenced (GATC/Eurofins). The sequence encoding the MANEA-Δ97 E404Q variant was generated by mutagenesis of the pColdI-MANEA construct using Q5 Site-Directed Mutagenesis Kit (NEB).

#### Truncated MANEA expression and protein purification

The *Homo sapiens MANEA-Δ97* gene was co-overexpressed with *groEL* and *groES* genes encoding the GroEL chaperone using a modified version of a method described previously (2). BL21(DE3) competent cells (Agilent) harboring pGro7 vector (TaKaRa Bio, Inc.) were transformed with pColdI-*MANEA-Δ97* and used to inoculate an LB starter culture containing 20 μg/ml chloramphenicol and 100 μg/ml ampicillin. The culture was shaken overnight at 37 °C in a 50 ml Falcon tube. The fully-grown starter culture was used to inoculate (1:100 v/v) the media for large-scale expression. The expression media was TB with 20 mM $MgCl_2$ added in order to promote the stability of the GroEL chaperone, 0.5 g/l of L-arabinose, 20 μg/ml chloramphenicol and 100 μg/ml ampicillin; the cells were shaken until the $OD_{600}$ was at least 0.8. Expression of MANEA-Δ97 was induced by a 5 min cold shock in an ice-water slurry and incubation (with no shaking) for 30 min at 15 °C, and subsequent addition of IPTG up to a concentration of 0.2 mM. The cultures were then shaken overnight at 15 °C and the pellet was harvested by centrifugation of the cell culture.

MANEA-Δ97 and MANEA-Δ97-E404Q mutant were purified using nickel or cobalt affinity chromatography, followed by cation exchange chromatography using and SP column (GE). The cation exchange buffer was 50 mM potassium phosphate pH 7.0, 50 mM KCl and the chromatography was performed using a 50 → 500 mM KCl gradient. MANEA-Δ97 eluted at a

KCl concentration of 185-200 mм. The proteins were stable up to a concentration of 20 mg $ml^{-1}$ in 25 mм HEPES pH 7.0, 200 mм NaCl and in 50 mм potassium phosphate pH 7.0, 185 mм KCl.

**Crystallization of truncated MANEA**

Crystal structures of 5 crystal forms of wild-type MANEA-Δ97 were solved. All wild-type enzyme crystals were grown at 19 °C. For all crystallization experiments, the vapor diffusion sitting drop method was used and the plates were kept in darkness.

Crystal form 1 was obtained from a 300 nl droplet (150 nl protein solution, 150 nl reservoir solution) in 100 mм MIB buffer pH 6.0. The protein was kept in 50 mм potassium phosphate pH 7.0, 50 mм KCl buffer at 5.5 mg/ml.

Crystal form 2 was obtained in 1 м sodium succinate pH 7.0, 1% (w/v) PEG-MME2000. The droplet contained 100 nl of protein solution and 200 nl of the reservoir solution. The protein was kept in 25 mм HEPES pH 7.0, 200 mм NaCl buffer at 10 mg/ml.

Crystal form 3 was obtained from a 300 nl droplet (150:150 or 100:200 nl protein:reservoir ratio yielded crystals) with the reservoir being 100 mм HEPES pH 7.5, 200 mм $MgCl_2$, 30% v/v PEG 400 (Alpha Aesar), 1 mм TEW (Anderson–Evans polyoxotungstate $[TeW_6O_{24}]^{6-}$ (3)). The protein was kept in 25 mм HEPES pH 7.0, 200 mм NaCl buffer at 10 mg/ml. Importantly, only PEG 400 from Alpha Aesar was able to yield crystals.

Crystal form 4 was obtained from a 300 nl droplet (150:150 nl protein:reservoir ratio) in 100 mм bis-tris pH 5.5, 25% w/v PEG 3350. The protein was kept at 10 mg/ml in 25 mм HEPES pH 7.0, 200 mм NaCl buffer with 2.23 mм GlcIFG and 2.23 mм α-1,2-mannobiose (10 x molar ratio) added.

The MANEA-Δ97-E404Q crystal used for the data collection (crystal form 5) was obtained from a 500:500 nl droplet, where the reservoir solution was 100 mм sodium acetate pH 4.6, 200 mм ammonium sulfate, 12.8% w/v PEG-MME 2000. The protein was kept at 10 mg $ml^{-1}$ in 25 mм HEPES pH 7.0, 200 mм NaCl buffer with 2.23 mм $GlcMan_4OMe$ (10 x molar ratio) added. The crystals were grown at 6 °C.

**Structure solution and refinement**

Diffraction data were collected at Diamond Light Source synchrotron at beamlines I03, I04 and I24. Crystallographic data were indexed and integrated using Diamond Light Source auto-processing pipelines that incorporate DIALS (4) software into Xia2 (5) or in-house using Xia2.

Data truncation, merging and scaling was performed in AIMLESS (6), **Supplemental Table S1, S2**. In case of two datasets (see **Supplemental Table S3**), the data were cut anisotropically using STARANISO software (7) in order to improve electron density maps and remove artefacts. Crystal form 1, whose structure was solved first, was solved using molecular replacement (MR) in PHASER (8) with the initial model being the *Bx*GH99 from PDB 5M17, truncated in CHAINSAW (9) and in COOT (10). Subsequent crystal forms were solved using truncated versions of the final model from crystal form 1 and crystal form 2 (chain A only). Only the polypeptide chain was used for the MR. Subsequently, the structures were refined in REFMAC5 (11) and real-space refinement was done using COOT. In case of isomorphous crystals in complex with different ligands, the HKL index of the new complex was matched to the first obtained solution, the $R_{free}$ set was copied and a model containing the protein only was directly refined to the observed data. Waters were added after the refinement of the polypeptide chain was complete and the ligand molecules were added in the last steps of the refinement. The model geometry and the correspondence of the model to experimental data were validated using COOT validation tools and the PDB validation pipeline. Sugars and pseudosugars were validated using PRIVATEER (12, 13). The process of model building, refinement and validation was performed using the CCP4i2 GUI (14).

The structure of *Bx*GH99 was solved using PDB 5M5D (protein only) as the starting model using the same protocol as in (15).

**Enzymology**

Activity of MANEA-Δ97 on $GlcMan_9GlcNAc_2$ was demonstrated by detection of the product by mass spectrometry. MANEA-Δ97 (100 nM) was incubated with 24.5 μM $GlcMan_9GlcNAc_2$ for 19 h at 4 °C, 14 h at RT and then 12.5 h at 37 °C in 50 mM potassium phosphate pH 7.0, 185 mM KCl buffer. The volume of the mixture was 20 μl. A negative control containing the substrate only was subjected to the same treatment. Subsequently, the sugars were permethylated according to the protocol described by Ciucanu and Kerek (16). Analysis of the content of the mixture was performed using MALDI-MS in the University of York Technology Facility by Dr Adam Dowle.

Michaelis-Menten kinetics were performed using a previously published method (17) where the substrate was $GlcMan_3OMe$ instead of $Man_4OMe$. The $K_M$ and $IC_{50}$ values (for inhibitors) were determined using ORIGIN (18). The concentration of the enzyme for all kinetics experiments was 1 μM. The buffer for the determination of $K_M$ was 50 mM potassium phosphate, pH 7.0, 185 mM KCl. HEPES buffers (for the determination of HEPES $IC_{50}$) contained 100–1800 mM of HEPES pH 7.0, 200 mM NaCl. Higher concentrations of HEPES were not possible due to the limited solubility of the buffer.

**Thermal shift assay**

Thermal unfolding was assessed using an Agilent Technologies Mx3005P qPCR system. MANEA-Δ97 was dissolved in 50 mM potassium phosphate pH 7.0, 185 mM KCl buffer. The final concentration of MANEA-Δ97 was 1 mg/ml. The temperature was increased from 25 to 95 °C in 1-degree increments of 30 s each. The reporter molecule was SYPRO Orange (fluorescence measured at 570 nm). The reporter was used at a final concentration of 2.5 × (1/2000 of the 5000× concentrate stock). The measurement was conducted in triplicate with two buffer controls.

**Circular dichroism**

Circular dichroism spectra were recorded using a Jasco J810 CD Spectrophotometer with a prism monochromator. The samples were kept in a 1 mm quartz cuvette. MANEA-Δ97 at 0.2 mg/ml was stable in low salt conditions (2.2 mM KCl). Ellipticity data were collected every 0.5 nm for wavelengths from 190 to 260 nm. Before data analysis, the buffer signal was subtracted from the sample signals. Data were analyzed using the CAPITO web server (19); the CD signal at a wavelength of 222 nm is produced by from alpha-helical structures.

**Isothermal Titration Calorimetry**

Isothermal titration calorimetry was performed using a MicroCal Auto-iTC200 (GE/Malvern Instruments). The protein was present in the cell and inhibitors in the syringe. Care was taken to perform experiments at a c-value of 5–1000 (c=[Protein]/$K_d$ (dimensionless)). The $K_d$ values, molar ratio, T$\Delta S$ and $\Delta H$ were calculated using a modified version of Origin7 program with ITC200 adjustments or using MicroCal PEAQ-ITC Analysis Software (Malvern Instruments).

*Bt*GH99 (*Bacteroides thetaiotaomicron* GH99) binding assays were carried out in triplicate at 25 °C, with 18 × 2 µl injections of CyMe-GlcIFG (500 µM) titrated into the ITC cell containing 50 µM *Bt*GH99 in 25 mM HEPES pH 8.0, 50 mM NaCl.

MANEA-Δ97 isothermal titration calorimetry was done at 25 °C in 50 mM potassium phosphate pH 7.0, 185 mM KCl buffer. Results of initial experiments suggested the number of ManIFG and GlcIFG binding sites (N) was ~2.3. To exclude the possibility of spurious binding, parallel experiments with *Bt*GH99 and MANEA-Δ97 were run to calibrate the ligand concentration. This reduced the discrepancy to ~1.7. Further calibration was achieved by adding 2 a saturating amount of α-1,2-mannobiose (1 mᴍ) to the cell and the syringe; the N reduced further to ~1.1. The calculated $K_d$, Δ*H* and -TΔ*S* were similar to those calculated from the experiments without α-1,2-mannobiose with N fixed at 1. We therefore assumed that the higher number of sites is an artefact and used N fixed at 1, letting the ligand concentration refine. The ITC experiments in the presence of α-1,2-mannobiose were performed with 44 µᴍ of MANEA-Δ97 in cell, 608 µᴍ of GlcIFG in syringe and 1 mᴍ of α-1,2-mannobiose in both. To exclude the possibility of protein concentration being wrong, MANEA-Δ97 in the sample after the ITC procedure was double-checked by unfolding the protein in 6 ᴍ guanidinium hydrochloride to expose all aromatic residues and measuring $Abs_{280}$ using an Eppendorf BioPhotometer (Mw=44719 kDa, ε=98670 $ᴍ^{-1}$ $cm^{-1}$). A single experiment at an optimal concentration of the inhibitor was selected and shown; error values are as reported by the MicroCal PEAQ-ITC Analysis Software.

**Cell culture and Virus Stock**

Mardin-Darby Bovine Kidney (MDBK) cells (European Collection of Animal Cell Cultures, Porton Down, United Kingdom) were cultured in RPMI-1640 medium (PAA Laboratories, Teddington, UK) supplemented with 10% fetal calf serum (PAA), 2 mᴍ ʟ-glutamine (PAA) and 100U/mL penicillin–streptomycin (PAA), at 37 °C in a humidified atmosphere containing 5% $CO_2$ (v/v). The human hepatoma cell line Huh7.5 (Apath, LLC) was cultured in Dulbecco's modified Eagle medium (DMEM) with Glutamax (Gibco) supplemented with 100 µg $mL^{-1}$ streptomycin (Sigma), 100 U $mL^{-1}$ penicillin (Sigma), 100 µm MEM non-essential amino acids (Gibco) and 10% (v/v) heat-inactivated (56 °C, 30 min) fetal calf serum (FCS, Seralab). For *in vitro* infection assays, BVDV-1 (cytopathic NADL strain) was provided by Professor Nicole Zitzmann (Oxford Glycobiology Institute, University of Oxford, UK). DENV2 strain 16681 (a gift from Andrew Webb, Screaton Lab (Imperial college London, UK) was propagated in C6/36 *Aedes albopictus* cell line (US Armed Forces Research Institute of Medical Sciences, Thailand (AFRIMS)), collected from supernatant, and concentrated by precipitation with 10% weight per unit volume (w/v) poly(ethyleneglycol) $M_r$ 6,000 (Sigma), 0.6% sodium chloride (Sigma) overnight at 4 °C. Following precipitation, virus was centrifuged at 2830 *g* for 45 min at 4 °C, resuspended in Leibovitz's L15 + 10% HI-FBS, and stored at –80 °C until use.

**Fluorescence focus assay**

MDBK cells were seeded in 6-well plates (1.75 × $10^6$ cells/well) and infected with BVDV at an MOI of 1, followed by incubation with different concentrations of GlcIFG (0-1 mᴍ) and in combination with NAP-DNJ (*N*-(6'-[4″-azido-2″-nitrophenylamino]hexyl)-1-DNJ; synthesised in house, Oxford Glycobiology Institute) for 24 h. Cell culture medium was then harvested and clarified by centrifugation at 5000 rpm for 10 min to remove cell debris. Serial dilutions were made and used to infect naive MDBK cells seeded in 96-well plates ($7 \times 10^4$ cells/well). Cells were incubated for 24 h, then washed twice with cold PBS and fixed with 4% (v/v) paraformaldehyde in PBS for 30 min. Cells were washed again and blocked for two hours in 5% (w/v) milk in PBS. Cells were then permeabilized with 1% (v/v) Triton-X-100 in PBS for 20 min, washed with 1% (v/v) Tween 20 in PBS and incubated with MAb103/105 (1:500 dilution in 5% milk-PBS; Animal Health Veterinary Laboratories Agency, Weybridge, UK) for 1 h. Cells were washed three times with 1% Tween 20-PBS then incubated with anti-mouse FITC-conjugated secondary antibody (1:500 dilution in 5% milk-PBS; Sigma). Cells were washed again, and nuclei stained with 4′,6-diamidino-2-phenylindole (DAPI; Vector Laboratories Inc.). Cells were observed and evaluated using an inverted Nikon Eclipse TE200-U microscope, equipped with a Fluor 10x objective lens, and FITC and UV filter sets. Fluorescent foci were counted, and virus titers (fluorescent focus units (FFU/ml) calculated.

**Immunoblotting**

MDBK cells were seeded in 6-well plates ($1.75 \times 10^6$ cells/well) and infected with BVDV at an multiplicity of infection (MOI) of 1, followed by incubation with different combinations of GlcIFG (1 mM) and 1 μM NAP-DNJ for 24 h. Cells were harvested by scraping in PBS and centrifuged at 2000 rpm for 5 min. The resulting pellet was briefly frozen at −20 °C then resuspended in PBS. The suspension was sonicated on ice using a Branson Sonifier (20 s, 50% duty cycle) to lyse cells. Protein concentrations were determined using a standard bicinchoninic acid protein assay. The samples were treated with Endo H (NEB, as per manufacturer’s protocol). Treated proteins were separated by sodium dodecyl sulfate–polyacrylamide gel electrophoresis (SDS–PAGE) under non-reducing conditions and transferred to Immobilon-P PVDF membrane (Millipore) by semi-dry electroblotting, then blocked overnight in 5% (w/v) milk, 0.2% (v/v) Tween 20 in PBS. The membrane was incubated with MAb214 primary antibody (1:1000 dilution in blocking solution; Animal Health Veterinary Laboratories Agency) for 2 h at room temperature, washed extensively with PBS–0.1% (v/v) Tween 20, then incubated with anti-mouse horseradish peroxidase secondary antibody (1:1000; Dako) for 1 h. Antibody binding to BVDV E2 protein was detected using Pierce ECL Western Blotting

Substrate (Thermo Scientific, UK) and X-ray film (GE Healthcare, UK) by following the manufacturer's instructions.

**DENV antiviral assay**

Huh7.5 cells were seeded in a 24-well plate at a density of $2 \times 10^5$ cells/well in 1 ml of growth media. After 18 h incubation, the supernatant was removed, and the wells inoculated for 2 h with 50 µL of DENV2 strain 16681 at a MOI of 1 in minimal essential media (MEM, Gibco). The inoculum was removed and replaced with 1 ml of growth media with drug at concentrations as indicated in triplicate, and the infected cells were incubated with drug for 2 d. The supernatant was harvested and stored at –80 °C until analysis by viral quantification or plaque assay.

**Viral RNA quantification**

DENV RNA in cell culture supernatants was isolated according to the manufacturer's protocol for Qiagen QIAamp Viral RNA Mini Kit and assayed by reverse transcription-real time polymerase chain reaction (qRT-PCR) on an Applied Biosystems 7500 real-time PCR system (Life Technologies). Thermo-Start DNA Taq polymerase was used to enable Taq-mediated release of fluorescently labelled dyes from a DENV2 NS5-specific probe adapted from a previously published protocol (20). Probe and primer sequence and concentrations were as previously described, with the reaction mixture prepared according to the manufacturer's instructions for Verso 1-Step RT-PCR Kit with Thermo-Start Taq (Life Technologies). Thermal cycling was adapted to match enzyme components as follows. Synthesis of complementary DNA (cDNA) was performed for 30 min at 50 °C, followed by a 15-min activation of Thermo-Start *Taq* polymerase at 95 °C. PCR thermocycling with fluorescence detection was executed for 45 cycles of 95 °C for 15 seconds followed by 60 °C for 60 s and a fluorescence read step. Samples were read in technical duplicate and compared to a standard curve generated from high-titer viral RNA isolated from C6/36-grown DENV2. 95% confidence intervals were determined based on biological and technical variation and graphed using Prism 6 (GraphPad Software, Inc.).

**DENV plaque assay**

The infectious DENV titres in collected supernatants were evaluated by plaque assay as described previously (21). Briefly, LLC-MK2 cells were seeded in confluent monolayers in a 12-well plate. Four 10-fold dilutions of each supernatant to be analyzed were prepared in

MEM. The adherent monolayers were washed once with Hank's buffered salt solution (Gibco), then inoculated with 100 µL of analyte and 100 µL MEM. The plates were rocked at 20 ℃ for 90 min, after which the inoculum was removed and nutrient-supplemented primary overlay (1 mL) was added to each well. The overlay was allowed to solidify at 20 °C, before returning the cells to the incubator. After 5 d, nutrient-supplemented, neutral red-containing secondary overlay (1 mL) was added to each well and allowed to solidify at 20 °C, then the cells were returned to the incubator. After 1 d, plaques were observed and counted from the underside of the plate by eye.

### Chemistry

#### General Methods

Thin layer chromatography (TLC) monitoring of reactions was performed using aluminium-backed plates of Merck Silica Gel 60 $F_{254}$ and detection was achieved using UV light or by charring with 5% sulfuric acid in methanol or staining with ceric ammonium molybdate (CAM; solution made from 1:5:10:90 $Ce(SO)_4/(NH_4)_6Mo_7O_{24}.4H_2O/H_2SO_4/H_2O$). Flash chromatography was performed using Geduran silica gel according to the method of Still *et al.*(22) Solvents were evaporated under reduced pressure using a rotary evaporator. Methanol was distilled over $CaH_2$. DMF was dried over activated 4 Å molecular sieves. Dichloromethane and diethyl ether were dried using a Grubbs solvent drying apparatus (Glass Contour of SG water, Nashua, U.S.A.).(23) Melting points were obtained using a Reichert–Jung hot stage melting point apparatus. Optical rotations were obtained using a JASCO DIP-1000 polarimeter (Melbourne, Australia). $[\alpha]_D$ values are given in deg.dm$^{-1}$ cm$^3$ g$^{-1}$. $^1$H, $^{13}$C and 2D NMR were recorded using a Varian Inova-500 (499.7 MHz for $^1$H and 125.8 MHz for $^{13}$C) at 25.0 °C. All signals were referenced to solvent peaks ($CDCl_3$: δ 7.26 ppm for $^1$H and 77.16 ppm for $^{13}$C; $D_2O$: δ 4.80 ppm for $^1$H). *J* values are given in Hz. A prime (') annotation for disaccharides specifies the carbohydrate ring at the non-reducing end; A, B, C, D is used for tetrasaccharides to indicate sugar rings from the non-reducing end; 'a' and 'b' indicate geminal protons. High resolution mass spectra (HRMS) were acquired on an Agilent ESI-TOF or Thermo Fisher Orbitrap.

#### Synthesis of Glc(Man)$_3$OMe

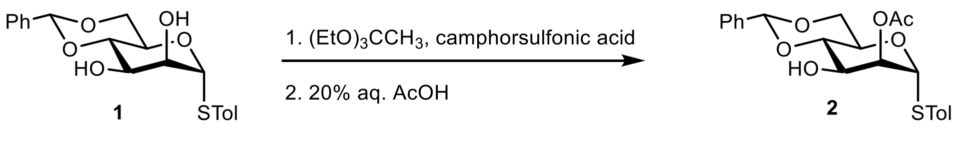

##### 4-Methylphenyl 2-*O*-acetyl-4,6-*O*-benzylidene-1-thio-α-D-mannopyranoside (2)

Camphorsulfonic acid (49 mg, 0.211 mmol) was added to a solution of 4-methylphenyl 4,6-*O*-benzylidene-1-thio-α-D-mannopyranoside(24) **1** (361 mg, 0.965 mmol) in triethyl orthoacetate (1.77 mL) and acetone (5.5 mL) and the reaction mixture was stirred at rt for 30 min. Acetic acid (20% aq.; 10.2 mL) was added to the cooled (0 °C) reaction mixture and stirred at rt for 1 h. The solvents were evaporated under reduced pressure and the resulting residue was subjected to flash chromatography (EtOAc/pet. spirits 35:65) to give the title alcohol (305 mg, 76%) as a colourless foam; $[\alpha]_D^{24}$ +158 (*c* 0.19, $CHCl_3$); $\delta_H$ (500 MHz, $CDCl_3$) 2.17 (3 H, s, Me), 2.34 (3 H, s, Me), 3.84 (1 H, apt. t, $J_{5,6a}$ = $J_{6a,6b}$ = 10.3 Hz, H6a), 3.99 (1 H, t, $J_{3,4}$ = $J_{4,5}$ = 9.7 Hz, H4), 4.24-4.28 (2 H, m, H3,6b), 4.38 (1 H, dt, $J_{4,5}$ = 9.9, $J_{5,6a}$ = 4.9 Hz, H5), 5.41 (1 H, d, $J_{1,2}$ = 1.0 Hz, H1), 5.49 (1 H, dd, $J_{1,2}$ = 3.6, $J_{2,3}$ = 1.3 Hz, H2), 5.62 (1 H, s, C**H**Ph), 7.13 (2 H, d, *J* = 7.9 Hz, Ar), 7.36-7.41 (5 H, m, Ar), 7.51-7.53 (2 H, m, Ar); $\delta_C$ (125 MHz, $CDCl_3$) 21.1 ($CH_3CO$), 21.3 (Ar**Me**), 64.7, 67.9, 68.6, 73.7, 79.3, 87.4, 102.4 (7 C, C1-C6,**C**HPh), 126.4, 128.5, 129.4, 129.5, 130.1, 132.9, 137.2, 138.5 (9 C, Ar, 3 × Cq), 170.5 (C=O).

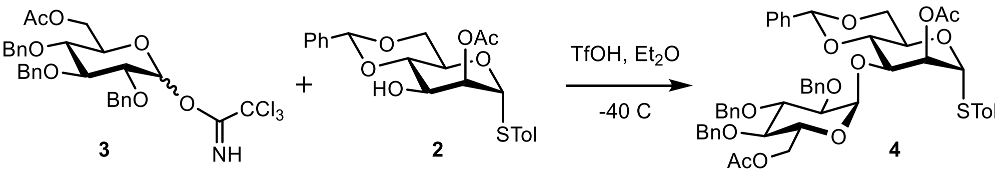

**4-Methylphenyl 2-*O*-acetyl-3-*O*-(6-*O*-acetyl-2,3,4-tri-*O*-benzyl-α-D-glucopyranosyl)-4,6-*O*-benzylidene-1-thio-α-D-mannopyranoside (4)**

TfOH in toluene (10% v/v, 60 µL, 0.068 mmol) was added to a stirred mixture of 6-*O*-acetyl-2,3,4-tri-*O*-benzyl-D-glucopyranosyl trichloroacetimidate(25) **3** (477 mg, 0.749 mmol), alcohol **2** (206.4 mg, 0.495 mmol) and freshly activated 4 Å molecular sieves in dry diethyl ether (25 mL) at -40 °C and the reaction mixture was stirred under $N_2$ for 30 min. The reaction was quenched with $Et_3N$ and the mixture was filtered through a Celite pad. The solvent was evaporated and the resulting residue was subjected to flash chromatography (EtOAc/pet. spirits 30:70 and $Et_2O/CHCl_3$ 0:100 to 1:100) to give the title disaccharide (180.7 mg, 41%) as a colourless oil; $[\alpha]_D^{23}$ +120 (*c* 0.53, $CHCl_3$); $\delta_H$ (500 MHz, $CDCl_3$) 2.10 (3H, s, Ac), 2.19 (3 H, s, Ac), 2.34 (3 H, s, Ar**Me**), 3.36 (1 H, dd, $J_{3,4}$ = 9.0, $J_{4,5}$ = 10.0 Hz, H4), 3.45 (1 H, dd, $J_{1,2}$ = 3.7, $J_{2,3}$ = 9.7 Hz, H2), 3.85 (2 H, m, H3,6b'), 3.94 (1 H, m, H5), 4.17-4.38 (6 H, m, H6a,6b,6a',3',4',C**H**$_2$Ph), 4.45 (1 H, ddd, $J_{4',5'}$ = 9.9, $J_{5',6a'}$ = 9.9, $J_{5',6b'}$ = 4.9 Hz, H5'), 4.52 (1 H, *J* = 12.5 Hz, C**H**$_2$Ph), 4.56 (1 H, *J* = 11.2 Hz, C**H**$_2$Ph), 4.76 (1 H, *J* = 10.9 Hz, C**H**$_2$Ph), 4.85 (1 H, *J* = 11.2 Hz, C**H**$_2$Ph), 4.95 (1 H, *J* = 10.9 Hz, C**H**$_2$Ph), 5.33 (1 H, d, *J* = 3.7 Hz, H1), 5.42 (1 H, d, *J* = 1.1 Hz, H1'), 5.52 (1 H, broad s, C**H**Ph), 5.53 (1 H, dd, $J_{1',2'}$ = 1.1, $J_{2',3'}$ = 3.5 Hz, H2'), 6.95 (2 H, m, Ar), 7.12-7.43 (22 H, m, Ar); $\delta_C$ (125 MHz, $CDCl_3$) 21.0, 21.1 (2 C, 2 × **C**$H_3$CO), 21.2 (1 C, Ar**Me**), 63.5 (1 C, C6), 64.9 (1 C, C5'), 68.5 (1 C, C6'), 69.4 (1 C, C5), 70.7 (1 C, **C**$H_2$Ph), 71.6 (1 C, C3'), 73.4 (1 C, **C**$H_2$Ph), 74.7 (1 C, **C**$H_2$Ph), 75.5 (1 C, **C**$H_2$Ph), 77.1 (1 C, C4), 78.6 (1 C, C2'), 79.2 (1 C, C4'), 81.1 (1 C, C3), 87.5 (1 C, C1'), 97.3 (1 C, C1),

102.6 (1 C, C2), 126.4-132.7 (26 C, Ar), 137.1, 138.0, 138.5, 138.7 (4 C, Cq), 170.0, 170.9 (2 C, 2 × C=O); HRMS (ESI)$^+$ *m/z* 913.3228 [$C_{14}H_{22}N_2O_9$ (M+Na)$^+$ requires 913.3228].

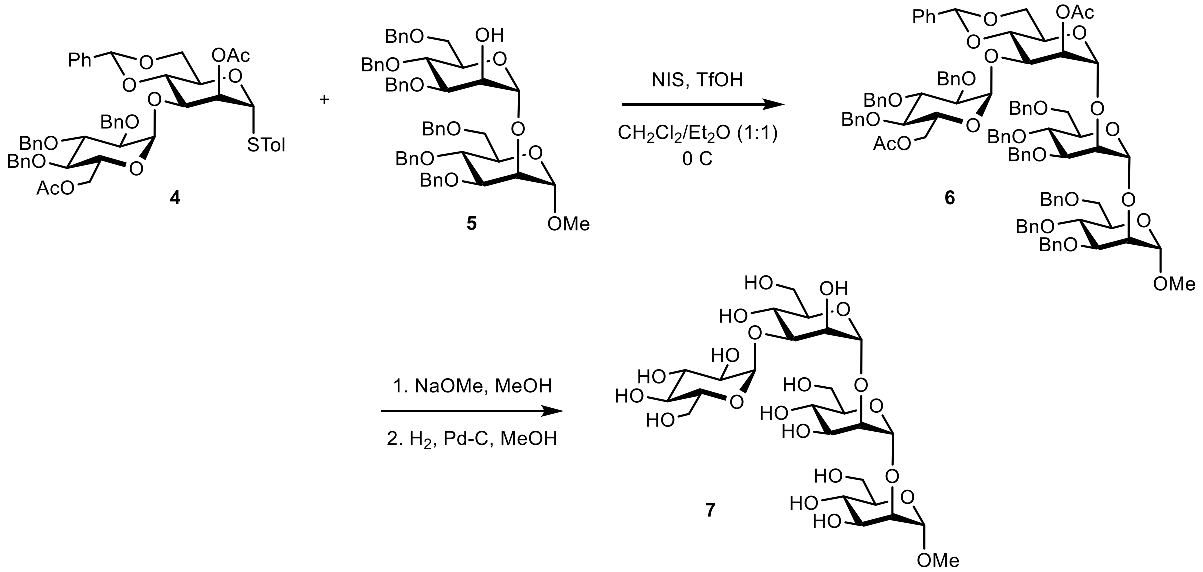

**Methyl 6-*O*-acetyl-2,3,4-tri-*O*-benzyl-α-D-glucopyranosyl-(1,3)-2-*O*-acetyl-4,6-*O*-benzylidene-α-D-mannopyranosyl-(1,2)-3,4,6-tri-*O*-benzyl-α-D-mannopyranosyl-(1,2)-3,4,6-tri-*O*-benzyl-α-D-mannopyranoside (6)**

NIS (10.5 mg, 0.05 mmol) and triflic acid (2.8 µL, 0.032 mmol) was added to a stirred suspension of the disaccharide donor **4** (52.1 mg, 0.058 mmol), methyl 2-*O*-(3,4,6-tri-*O*-benzyl-α-D-mannopyranosyl)-3,4,6-tri-*O*-benzyl-α-D-mannopyranoside(26) **5** (35.5 mg, 0.04 mmol) and freshly activated 4 Å molecular sieves in dry DCM/ $Et_2O$ (1:1, 1 mL) at 0 °C and the reaction mixture was stirred for 0.5 h. The reaction was neutralized with $Et_3N$, filtered through a Celite pad, the solvent was removed under reduced pressure and the resulting residue was subjected to flash chromatography (EtOAc/pet. spirits 30:70) to give the title tetrasaccharide (19 mg, 29%) as a colourless oil; $[\alpha]_D^{26}$ +49 (*c* 0.285, $CHCl_3$); $\delta_H$ (500 MHz, $CDCl_3$) 1.87 (3 H, s, Ac), 2.18 (3 H, s, Ac), 3.19 (3 H, s, Me), 3.44 (2 H, m, H2$^C$), 3.67-4.15 (17 H, m, H2$^B$,4$^D$), 4.27 (2 H, m), 4.40 (1 H, dd, $J_{2,3}$ = 3.6, $J_{3,4}$ = 9.9 Hz, H3$^D$), 4.48-4.85 (18 H, m, H1a,17 × C**H**$_2$Ph), 4.95 (1 H, d, *J* =10.8 Hz, C**H**$_2$Ph), 5.04 (1 H, d, $J_{1,2}$ = 1.1 Hz, H1$^D$), 5.17 (1 H, d, $J_{1,2}$ = 1.4 Hz, H1$^B$), 5.34 (1 H, d, $J_{1,2}$ = 3.6 Hz, H1$^C$), 5.42 (1 H, dd, $J_{1,2}$ = 1.4, $J_{2,3}$ = 3.5 Hz, H2$^D$), 5.44 (1 H, s, C**H**Ph), 6.96 (2 H, apt. d, Ar), 7.12-7.37 (48 H, Ar); $\delta_C$ (125 MHz, $CDCl_3$) 21.1 (2 C, 2 × **C**$H_3$CO), 54.7 (1 C, OMe), 60.4, 62.6, 64.1, 68.6, 69.1, 69.3, 69.5, 70.7, 71.0, 71.6, 71.7, 72.1, 72.2, 72.4, 73.3, 73.3, 74.6, 74.9, 75.0, 75.1, 75.3, 75.5, 76.5, 78.8, 79.2, 81.2 (27 C), 97.1 (1 C, C1$^C$), 99.7 (1 C, C1$^A$), 99.9 (1 C, C1$^D$), 100.8 (1 C, C1$^B$), 102.5 (1 C,

**C**HPh), 126.5-138.7 (50 C, Ar, 10 × Cq), 169.7, 170.4 (2 C, 2 × C=O); HRMS (ESI)$^+$ *m/z* 1675.7014 [$C_{14}H_{22}N_2O_9$ (M+Na)$^+$ requires 1675.7017].

**Methyl α-D-glucopyranosyl-(1,3)-α-D-mannopyranosyl-(1,2)-α-D-mannopyranosyl-(1,2)-α-D-mannopyranoside (7)**

Sodium methoxide in methanol (0.087 M, 0.42 mL, 0.036 mmol) was added to a solution of the diacetate **6** (19 mg, 0.01 mmol) in distilled methanol (2.7 mL) and the mixture was stirred overnight at 30 °C. The mixture was neutralized using Dowex-50 resin ($H^+$ form), filtered and the solvent evaporated to give a diol (16 mg) as a white residue. Pd/C (5%, 18 mg) was added to a solution of the crude diol in methanol (2 mL) and the reaction mixture was stirred under a hydrogen atmosphere (30 mbar) for 5 h. The suspension was filtered and the solids were washed with methanol. Reverse phase chromatography ($H_2O$, 100%) afforded the title tetrasaccharide (6.7 mg, 98% over two steps) as a colourless residue; $[\alpha]_D^{22}$ +65 (*c* 0.35, $CH_3OH$) (lit.(27) +101, $H_2O$); $\delta_H$ (500 MHz, $D_2O$) 3.43 (3 H, s, Me), 3.58-4.00 (22 H, m), 4.15 (1 H, br s, H2), 4.27 (1 H, br s, H2), 5.03 (1 H, br s, H1), 5.07 (1 H, br s, H1), 5.29 (1 H, d, $J_{1,2}$ = 3.8 Hz, H1), 5.33 (1 H, br s, H1); $\delta_C$ (125 MHz, $D_2O$) 53.7 (1 C, Me), 59.6, 59.8, 59.8, 60.0 (4 C, C6), 65.0, 65.8, 66.0, 68.6, 68.7, 68.8, 69.1, 70.7, 71.2, 71.4, 71.7, 72.2, 72.2, 77.2 (14 C), 77.4 (1 C, C2), 77.6 (1 C, C2), 98.2, 99.3, 99.5, 101.0 (4 C, C1); HRMS (ESI)$^+$ *m/z* 703.2269 [$C_{14}H_{22}N_2O_9$ (M+Na)$^+$ requires 703.2267].

**Synthesis of 3-*O*-cyclohexylmethyl-α-Glc-1,3-isofagomine (CyMe-GlcIFG)**

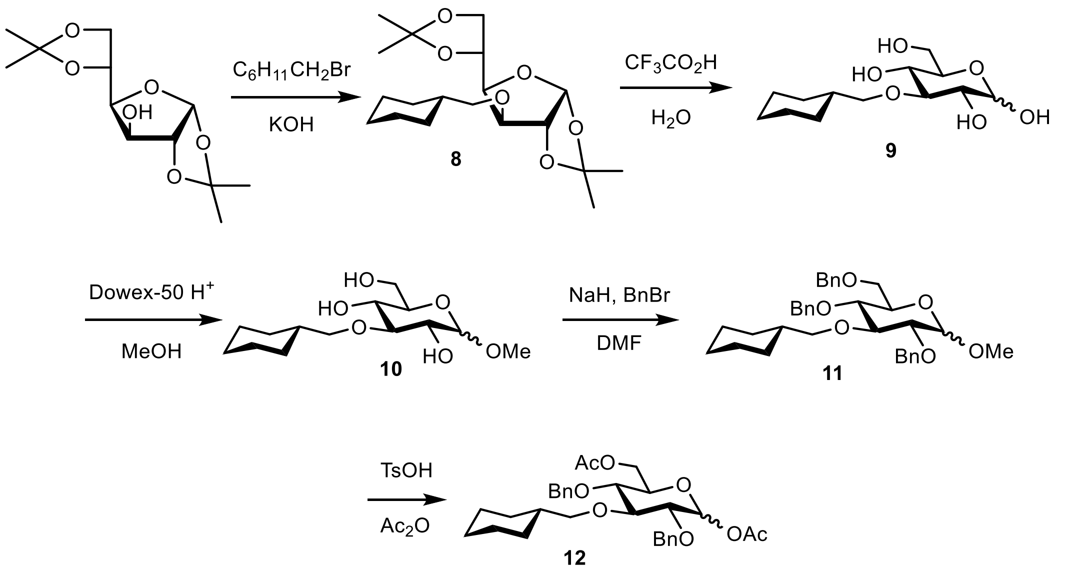

**3-*O*-Cyclohexylmethyl-1,2:5,6-di-*O*-isopropylidene-α-D-glucofuranose (8)**

Crushed potassium hydroxide (0.757 g, 13.5 mmol) was added to a stirred suspension of diacetone-D-glucose (0.260 g, 1.00 mmol) and cyclohexylmethyl bromide (0.488 mL, 3.50 mmol). The temperature was raised to 120 ºC and the reaction was stirred overnight. The mixture was cooled, diluted with water and extracted with chloroform (3 × 15 mL). The combined organic extracts were dried (MgSO4) and concentrated under reduced to yield a residue which was purified by flash chromatography (EtOAc/pet. spirits 0:100 to 10:90 + 1% TEA) to afford the title ether as a pale yellow oil (0.342 g, 96%), $[\alpha]_D^{24}$ -30 (*c* 0.285, $CH_2Cl_2$); $\delta_H$ (500 MHz, $CDCl_3$) 0.94, 1.21, 1.71 (11 H, 3 × m, $C_6H_{11}$) 1.31, 1.34, 1.42, 1.49 (12 H, 4 × s, Me), 3.32, 3.39 (2 H, ABq, $J_{H,H}$ = 9.1 Hz, $OCH_2$), 3.83 (1 H, d, $J_{3,4}$ = 3.1 Hz, H3), 3.97 (1 H, dd, $J_{5,6}$ = 6.3, $J_{6,6}$ = 8.5 Hz, H6), 4.08 (1 H, dd, $J_{5,6}$ = 6.3, $J_{6,6}$ =8.5 Hz, H6), 4.13 (1 H, dd, $J_{3,4}$ = 3.1, $J_{4,5}$ =7.4 Hz, H4), 4.31 (1 H, dd, $J_{4,5}$ = 6.2, $J_{5,6}$ = 6.3 Hz, H5), 4.51 (1 H, d, $J_{1,2}$ = 3.7 Hz, H2), 5.87 (1 H, d, $J_{1,2}$ = 3.7 Hz, H1); $\delta_C$ (125 MHz, $CDCl_3$) 25.6, 26.0, 30.06, 30.12, 38.2 (6 C, $C_6H_{11}$), 26.4, 26.7, 26.9, 27.0 (4 C, 4 × Me), 67.4 (1 C, C6), 72.7 (1 C, C5), 76.5, 81.4 (2 C, C2, $OCH_2$), 82.4, 82.6 (2 C, C3,4), 105.5 (1 C, C1), 109.0, 111.9 (2 C, **C**$(Me)_2$); HRMS $(ESI)^+$ *m/z* 379.2091 [$C_{19}H_{32}O_6$ $(M + Na)^+$ requires 379.2091].

**3-*O*-Cyclohexylmethyl-α/β-D-glucopyranose (9)**

The ether **8** (0.342 g, 0.961 mmol) was treated with 9:1 TFA/$H_2O$ (2 mL) and stirred at rt for 1 h. The reaction mixture was concentrated under reduced pressure and dried by azeotroping with toluene several times to remove volatile components. Flash chromatography

(EtOAc/MeOH/$H_2O$ 67:2:1) afforded the alkylated glucopyranose **9** as a white solid α:β = 0.03:1 (0.215 g, 81%), m.p. 126-133 ºC. NMR of major α-anomer: $\delta_H$ (500 MHz, $d_4$-MeOH) 1.00, 1.27, 1.72 (11 H, 3 × m, $C_6H_{11}$), 3.18 (2 H, m, $OCH_2$), 3.30 (1 H, m, H5), 3.37 (1 H, t, $J_{3,4}$ = 10.0 Hz, H3), 3.65 (3 H, m, H2,4,6), 3.87 (1 H, dd, $J_{6,6}$ = 2.0, 12 Hz, H6), 4.49 (1 H, d, $J_{1,2}$ = 7.5 Hz, H1); $\delta_C$ (125 MHz, $d_4$-MeOH) 27.0, 27.9, 31.2, 31.2, 39.9 (6 C, $C_6H_{11}$), 62.9 (1 C, C6), 71.5 (1 C, C4), 76.3 (1 C, C2), 78.0 (1 C, $OCH_2$), 80.0 (1 C, C5), 86.5 (1 C, C3), 98.3 (1 C, C1); HRMS $(ESI)^+$ *m/z* 299.1465 [$C_{13}H_{24}O_6$ $(M + Na)^+$ requires 299.1465].

**Methyl 3-*O*-cyclohexylmethyl-α/β-D-glucopyranoside (10)**

Cation exchange resin (Dowex 50W $H^+$ form; 0.500 g, resin pre-washed with water, 2 M HCl, water and MeOH) was added to a solution of 3-*O*-cyclohexyl-D-glucopyranose **9** (0.692 g, 2.51 mmol) in distilled MeOH (15 mL) and the mixture stirred overnight under reflux. The reaction was cooled to room temperature and the resin was removed by filtration. The filtrate was concentrated and purified by flash chromatography (EtOAc/MeOH/$H_2O$ 97:2:1) to afford the title compounds (α:β = 1.0:0.38) as a colourless oil (0.590 g, 81%). Partial $^1$H NMR: $\delta_H$ (500 MHz, $d_4$-MeOH) 1.00, 1.27, 1.70, 1.87 (11 H, 4 × m, α/β-$C_6H_{11}$), 3.45 (3 H, s, α-OMe), 3.57 (3H, s, β-OMe), 3.84 (1 H, dd, $J_{6,6}$ = 2.4, 11.8 Hz, α-H6), 3.90 (1 H, dd, $J_{6,6}$ = 2.3, 11.9 Hz, β-H6), 4.20 (1 H, d, $J_{1,2}$ = 7.5 Hz, β-H1), 4.67 (1 H, d, $J_{1,2}$ = 3.5 Hz, α-H1); $\delta_C$ (125 MHz, $d_4$-MeOH) 27.0, 27.9, 31.2, 39.90, 39.92 (6 C, α/β-$C_6H_{11}$), 57.5 (1 C, α-Me), 57.3 (1 C, β-Me), 62.6 (1 C, α-C6), 62.7 (1 C, β-C6), 71.4 (1 C, β-C4), 71.5 (1 C, α-C4), 73.6 (1 C, β-C2), 73.7 (1 C, α-C2), 75.1, 78.0 (2 C, α/β-$OCH_2$), 80.0 (1 C, β-C5), 80.2 (1 C, α-C5), 87.8 (1 C, α-C3), 86.5 (1 C, β-C3), 101.4 (1 C, α-C1), 105.5 (1 C, β-C1); HRMS $(ESI)^+$ *m/z* 314.1622 [$C_{14}H_{26}O_6$ $(M + Na)^+$ requires 314.1622].

**Methyl 2,4,6-tri-*O*-benzyl-3-*O*-cyclohexylmethyl-α/β-D-glucopyranoside (11)**

Methyl 3-*O*-cyclohexylmethyl-D-glucopyranosides **10** (0.170 g, 0.586 mmol) and NaH (70 mg, 1.76 mmol) were dissolved in DMF (3 mL) and stirred for 10 min. Benzyl bromide (210 µL, 2.34 mmol) was added dropwise at 0 ºC and the reaction warmed to rt. After 90 min NaH (17.5 mg, 0.439 mmol) was added and the mixture was stirred for 10 min. The reaction was cooled to 0 ºC and benzyl bromide (70 µL, 0.586 mmol) was added dropwise. This protocol was repeated once more after 90 min and the reaction mixture was stirred at rt overnight. The reaction was quenched with MeOH and water, and the mixture extracted with dichloromethane. The organic phase was washed with water (3 × 10 mL) and dried ($MgSO_4$), concentrated, and purified by flash chromatography (EtOAc/pet. spirits 0:100 to 10:90) to give the tribenzyl ether (α:β 1:0.6) as a colourless oil (0.319 g, 97%). NMR of the major α-anomer: $\delta_H$ (500 MHz, $CDCl_3$) 0.97, 1.20, 1.67, 1.81 (11 H, 4 × m, $C_6H_{11}$), 3.34 (3 H, s, OMe), 3.44 (1 H, dd, $J_{1,2}$ 3.6, $J_{2,3}$ = 9.6 Hz, H2), 3.52-3.75 (7 H, m, H3,4,5,6a,6b, $OCH_2$), 4.46, 4.83 (2 H, 2 × d, $J$ = 10.7 Hz, C**H**$_2$Ph), 4.47, 4.79 (2 H, 2 × d, $J$ = 12.2 Hz, C**H**$_2$Ph), 4.55 (1 H, d, $J_{1,2}$ = 3.6 Hz, H1), 4.62, 4.59 (2 H, 2 × d, $J$ = 11.8 Hz, C**H**$_2$Ph), 7.17-7.39 (15 H, m, 3 × Ph); $\delta_C$ (125 MHz, $CDCl_3$) 26.0, 26.8, 30.4, 30.5, 39.0 (6 C, $C_6H_{11}$), 55.2 (1 C, Me), 68.7 (1 C, C6), 70.1, 73.6, 73.7, 75.1, 77.9, 79.7, 80.0 (7 C, 3 × **C**H$_2$Ph, $OCH_2$, C2,4,5), 82.0 (1 C, C3), 98.5 (1 C, C1), 127.8-128.5 (15 C, 3 × Ph), 138.1, 138.5 (3 C, 3 × Ph); HRMS (ESI)$^+$ *m/z* 583.3031 [$C_{35}H_{44}O_6$ (M + Na)$^+$ requires 583.3030].

**1,6-Di-*O*-acetyl-2,4-di-*O*-benzyl-3-*O*-cyclohexylmethyl-α/β-D-glucopyranoside (12)**

TsOH.$H_2O$ (8.70 mg, 0.504 mmol) was added to a solution of the tribenzylated methyl glucopyranosides **11** (0.226 g, 0.403 mmol) in $Ac_2O$ (2.3 mL) and the mixture was stirred for 2 h at 70 ºC. The reaction was then poured into ice water and extracted with EtOAc. The organic phase was washed with water (2 × 10 mL), $NaHCO_3$ (2 × 10 mL), brine (10 mL) and dried ($MgSO_4$). The organic extract was concentrated and the residue was purified by flash chromatography (EtOAc/pet. spirits 15:85) to give the diacetates **12** (α:β 1:0.27) as a pale yellow oil (0.181 g, 83%), Partial $^1$H NMR: $\delta_H$ (500 MHz, $CDCl_3$) 0.97, 1.20, 1.70, 1.80 (11 H, 4 × m, $C_6H_{11}$), 2.03, 2.12 (6 H, 2 × s, α-Ac), 2.03, 2.04 (6 H, 2 × s, β-Ac), 5.57 (1 H, d, $J_{1,2}$ = 7.8 Hz, β-H1), 6.25 (1 H, d, $J_{1,2}$ = 3.6 Hz, α-H1), 7.27-7.38 (15 H, m, 3 × Ph); $\delta_C$ major α anomer: (125 MHz, $CDCl_3$) 21.0, 21.2 (2 C, 2 × Me), 26.0, 26.1, 26.7, 30.4, 30.5, 39.2 (6 C, $C_6H_{11}$), 62.9 (1 C, C6), 71.1, 73.5, 75.3, 79.0 (4 C, C5,$OCH_2$, 2 × **C**H$_2$Ph), 79.7 (1 C, C4), 81.7 (1 C, C3), 89.9 (1 C, C2), 104.9 (1 C, C1), 127.8-128.7 (10 C, 2 × Ph), 137.9, 137.8 (2 C, 2 × Ph), 169.5, 170.8 (2 C, 2 × C=O); HRMS (ESI)$^+$ *m/z* 563.2618 [$C_{31}H_{40}O_8$ (M + Na)$^+$ requires 563.2615].

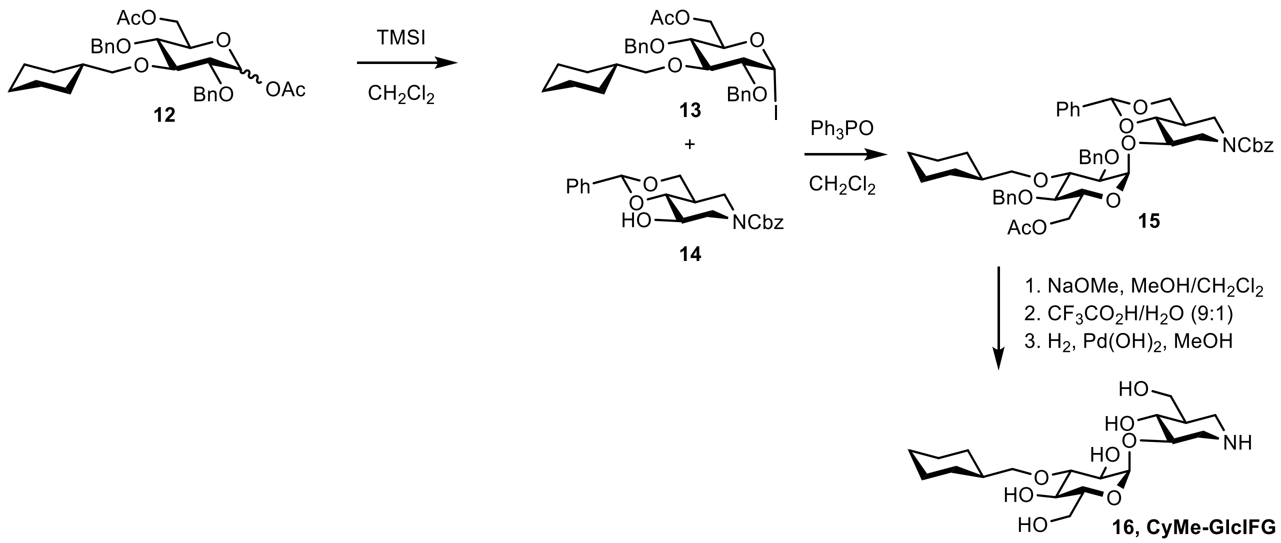

**3-*O*-(6'-*O*-Acetyl-2',4'-di-*O*-benzyl-3'-*O*-cyclohexylmethyl-α-D-glucopyranosyl)-(4,6-*O*-benzylidene-*N*-benzyloxycarbonyl-isofagomine) (15)**

TMSI (70.0 μL, 0.485 mmol) was added slowly to a solution of the diacetate **12** (0.175 g, 0.326 mmol) in dichloromethane (3 mL) at 0 °C and stirred for 10 min. Volatile materials were removed under reduced pressure by co-evaporation with distilled toluene under an atmosphere of nitrogen. Solutions of the crude glucosyl iodide **13** and acceptor(28) **14** (57.5 mg, 0.156 mmol), each in dichloromethane (5 mL) were added to a mixture of triphenylphosphine oxide (0.260 g, 0.934 mmol) and 4 Å molecular sieves and the mixture was stirred at rt for 7 d. The reaction mixture was filtered through Celite and washed with aq. $Na_2S_2O_3$ (3 × 5 mL) and brine (5 mL). Flash chromatography (EtOAc/pet. spirits 20:80 to 30:70) afforded the fully protected disaccharide **15** as a colourless oil (65.9 mg, 50%), $[\alpha]_D^{22}$ +64.7 (*c* 0.39, $CH_2Cl_2$); $\delta_H$ (500 MHz, $CDCl_3$) 0.97, 1.20, 1.68, 1.79 (11 H, 4 × m, $C_6H_{11}$), 2.02 (4 H, m, Ac, H5), 2.45, 2.87 (2 H, m, H2a,6a), 3.33 (1 H, dd, $J_{1',2'}$ = 3.7, $J_{2',3'}$ = 9.7 Hz, H2'), 3.36-5.16 (17 H, m, H2b,3,4,6b,7,7,3',4',5',6',6', 3 × $OCH_2$), 5.39 (1 H, d, $J_{1',2'}$ = 3.7 Hz, H1'), 5.54 (1 H, s, PhC**H**), 6.92 (2 H, m, PhC**H**$_2$), 7.13-7.41 (20 H, m, 4 × Ph); $\delta_C$ (125 MHz, $CDCl_3$) 20.9 (1 C, $CH_3$), 26.03, 26.05, 26.7, 30.3, 30.5, 37.7, 39.1 (7 C, $C_6H_{11}$, C5), 43.2 (1 C, C6), 47.2 (1 C, C2), 63.1, 67.8, 68.4, 69.2, 71.2, 71.6, 75.3, 76.8, 78.5, 79.7, 81.4, 84.9 (12 C, C3,4,7,2',3',4',5',6', 3 × **C**$H_2$Ph, $OCH_2$), 96.8 (1 C, **C**HPh), 102.5 (1 C, C1'), 126.6-129.5 (20 C, 4 × Ph), 136.3-137.9 (4 C, 4 × Ph), 155.0 (1 C, NC=O), 170.8 (1 C, C=$OCH_3$); HRMS $(ESI)^+$ *m/z* 872.3983 [$C_{50}H_{59}NO_{11}$ $(M + Na)^+$ requires 872.3980].

**(3*R*,4*R*,5*R*)-3-(3'-*O*-Cyclohexylmethyl-α-D-glucopyranosyloxy)-4-hydroxy-5-(hydroxymethyl)-piperidine (16, CyMe-GlcIFG)**

Sodium methoxide solution (70.0 μL, 0.071 mmol, 1.0 M) was added to the pseudo-disaccharide **15** (60.0 mg, 0.071 mmol) in $CH_2Cl_2$/MeOH (1:4, 3 mL) and stirred until TLC indicated conversion to a single compound of higher polarity (1 h). After neutralization with Dowex-50 resin ($H^+$ form), the mixture was filtered and concentrated. The residue was chromatographed (EtOAc/pet. spirits 35:65) and concentrated. The residue was treated with TFA/$H_2O$ (9:1) stirred at rt for 10 min (3 × 1 mL) and concentrated. The presumed triol was dissolved in MeOH (15 mL), treated with HCl (0.5 M, 1 mL), $Pd(OH)_2$/C (20%, 200 mg) then purged with hydrogen and left to stir under an atmosphere of hydrogen (18 h). The reaction mixture was filtered through a thick layer of Celite (washing thoroughly with 50% aq. MeOH) and concentrated under reduced pressure. The residue was purified by ion-exchange chromatography (Dowex 1X-8, $OH^-$ form, eluted with water; Dowex-50W-X2, $H^+$ form, eluted with water then 6 M aqueous $NH_3$) followed by reversed phase chromatography (MeOH/$H_2O$, 5:95) to afford CyMe-GlcIFG **16** as a colourless gum (19.7 mg, 69%), $[\alpha]_D^{23}$ + 79.3 (*c* 0.0975, EtOH); $\delta_H$ (500 MHz, $D_2O$/$d_4$-MeOH 1:1) 0.95, 1.26, 1.65, 1.79 (11 H, 4 × m, $C_6H_{11}$), 1.87 (1 H, m, H5), 2.70, 2.76 (2 H, m, H2,6), 3.28-3.85 (14 H, m, H2,3,4,6,7,2',3',4',5',6', $OCH_2$), 5.12 (1 H, d, $J_{1',2'}$ = 3.7 Hz, H1'); $\delta_C$ (125 MHz, $D_2O$/$d_4$-MeOH 1:1) 26.4, 27.3, 30.6, 30.7, 39.0 (6 C, $C_6H_{11}$), 43.9 (1 C, C6), 46.4 (1 C, C2), 48.4 (1 C, C5), 60.8 (1 C, C7), 61.8 (1 C, C6), 70.5, 72.3 (2 C, C4,4'), 72.7 (1 C, C2'), 73.8, 80.4, 81.0, 82.8 (4 C, $OCH_2$, C3,3',5'), 101.9 (1 C, C1'); HRMS $(ESI)^+$ *m/z* 406.2435 [$C_{19}H_{35}NO_8$ $(M + H)^+$ requires 406.2435].

### NMR spectra

##### 4-Methylphenyl 2-*O*-acetyl-4,6-*O*-benzylidene-1-thio-α-D-mannopyranoside (2)

$^{1}$H NMR (500 MHz, $CDCl_3$)

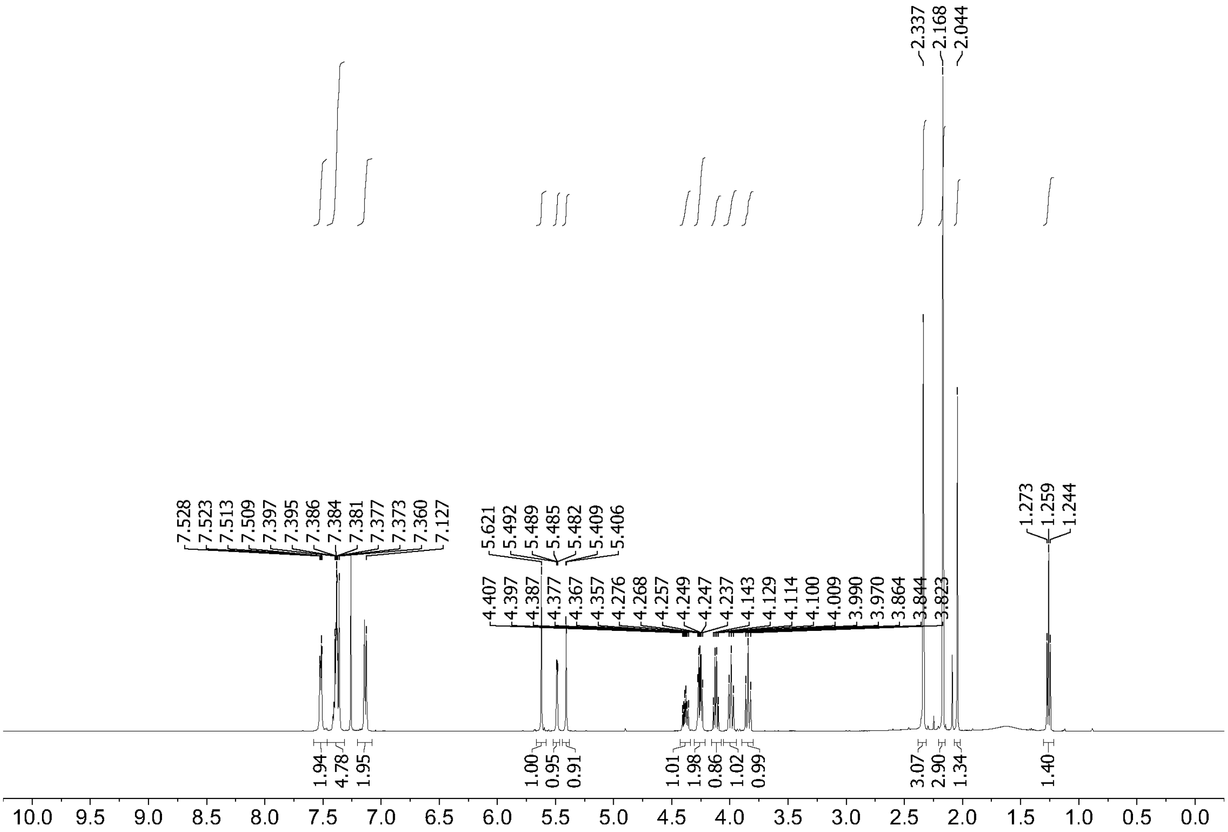

$^{13}$C NMR (125 MHz, $CDCl_3$)

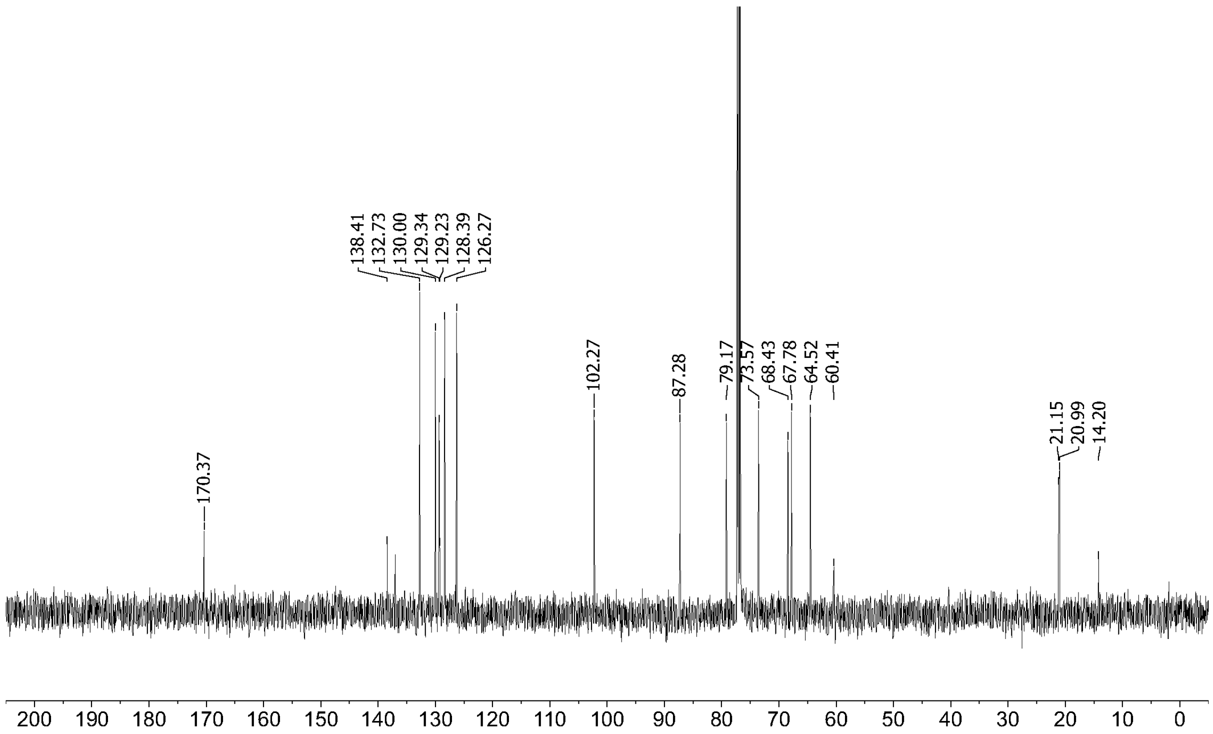
170.37
138.41
132.73
130.00
129.34
129.23
128.39
126.27
102.27
87.28
79.17
73.57
68.43
67.78
64.52
60.41
21.15
20.99
14.20
200 190 180 170 160 150 140 130 120 110 100 90 80 70 60 50 40 30 20 10 0

**4-Methylphenyl 2-*O*-acetyl-3-*O*-(6-*O*-acetyl-2,3,4-tri-*O*-benzyl-α-D-glucopyranosyl)-4,6-*O*-benzylidene-1-thio-α-D-mannopyranoside (4)**

$^{1}H$ NMR (500 MHz, $CDCl_3$)

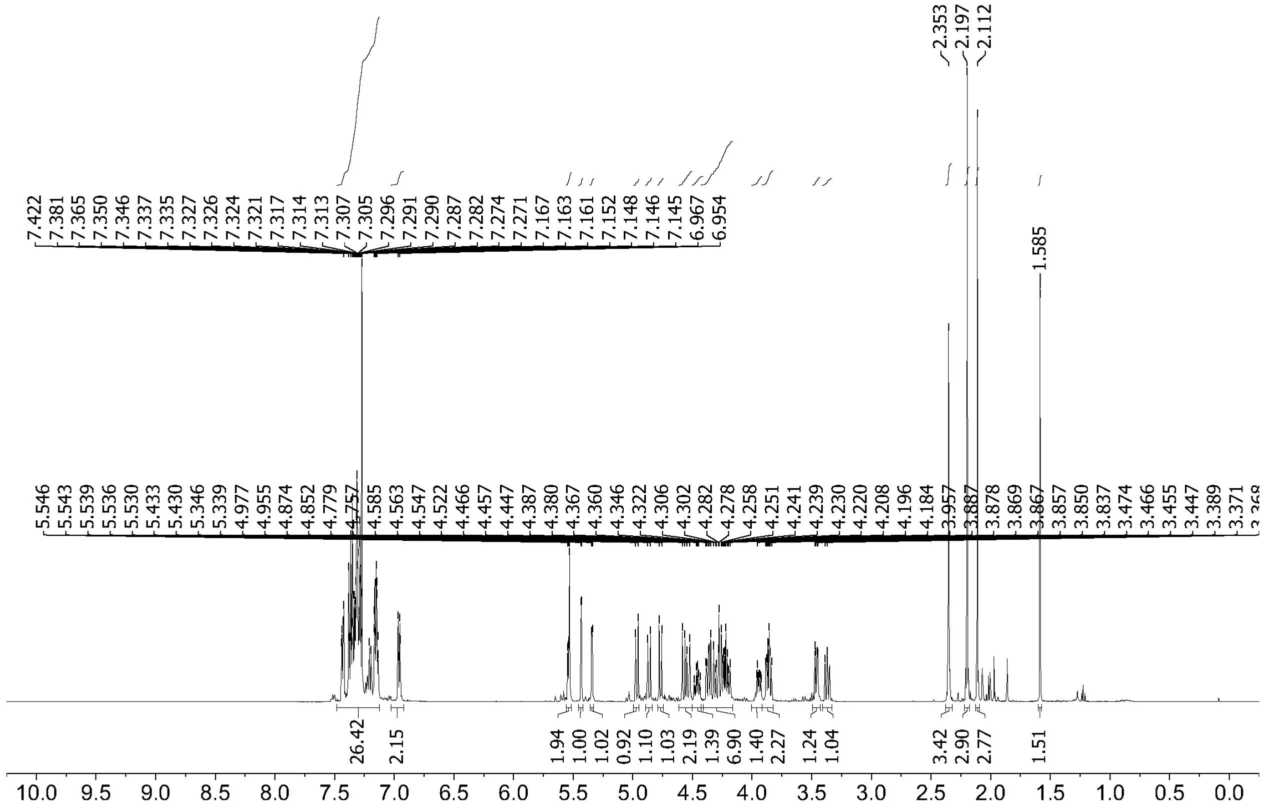

$^{13}C$ NMR (125 MHz, $CDCl_3$)

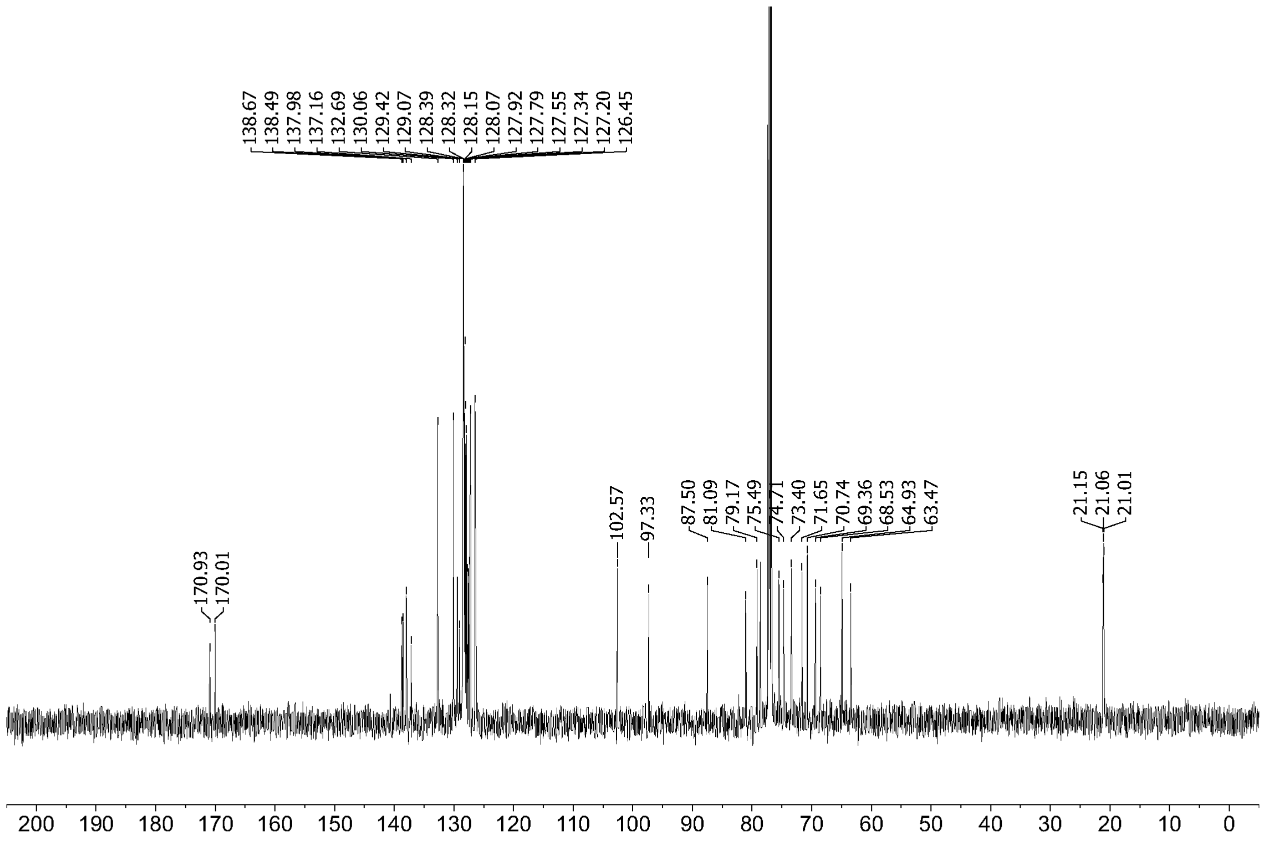
138.67
138.49
137.98
137.16
132.69
130.06
129.42
129.07
128.39
128.32
128.15
128.07
127.92
127.79
127.55
127.34
127.20
126.45
170.93
170.01
102.57
97.33
87.50
81.09
79.17
75.49
74.71
73.40
71.65
70.74
69.36
68.53
64.93
63.47
21.15
21.06
21.01
200 190 180 170 160 150 140 130 120 110 100 90 80 70 60 50 40 30 20 10 0

**Methyl 6-*O*-acetyl-2,3,4-tri-*O*-benzyl-α-D-glucopyranosyl-(1,3)-2-*O*-acetyl-4,6-*O*-benzylidene-α-D-mannopyranosyl-(1,2)-3,4,6-tri-*O*-benzyl-α-D-mannopyranosyl-(1,2)-3,4,6-tri-*O*-benzyl-α-D-mannopyranoside (6)**

$^1$H NMR (500 MHz, $CDCl_3$)

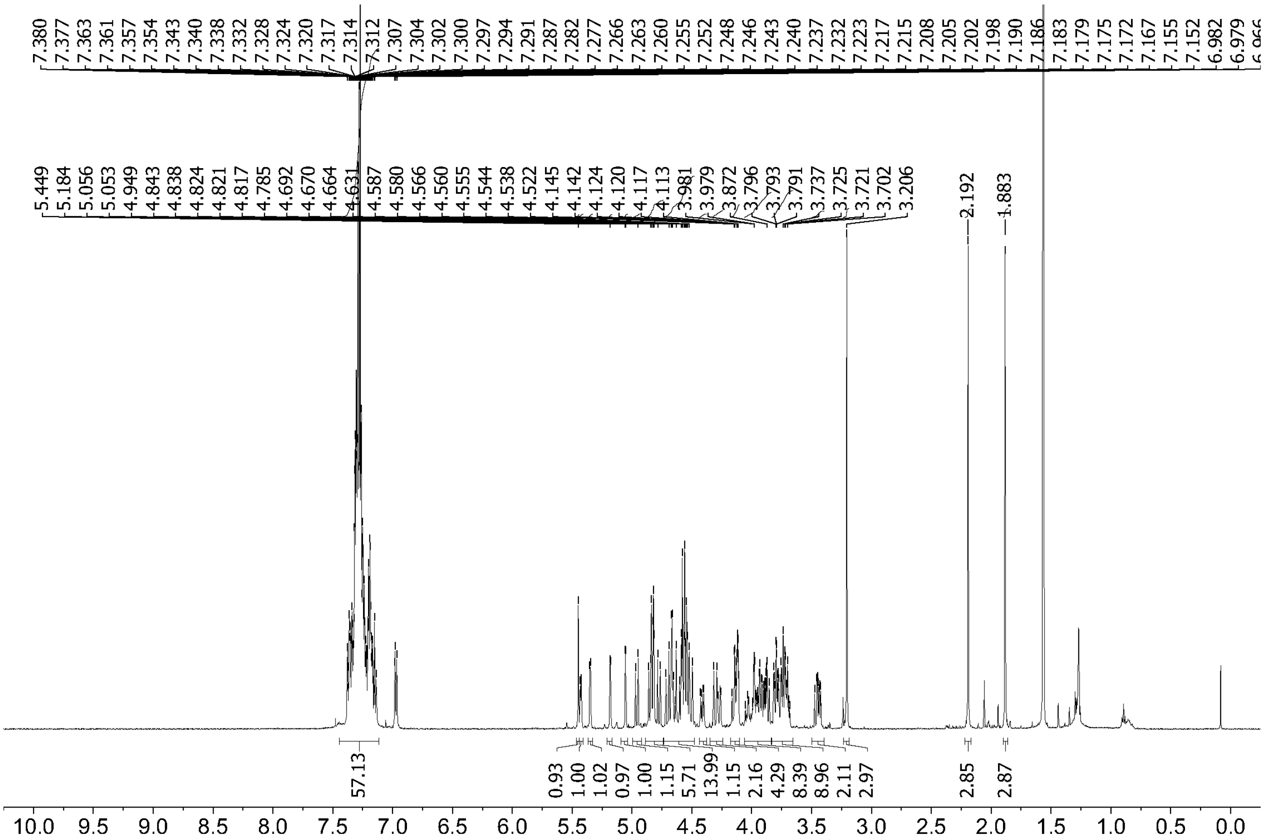

$^{13}$C NMR (125 MHz, $CDCl_3$)

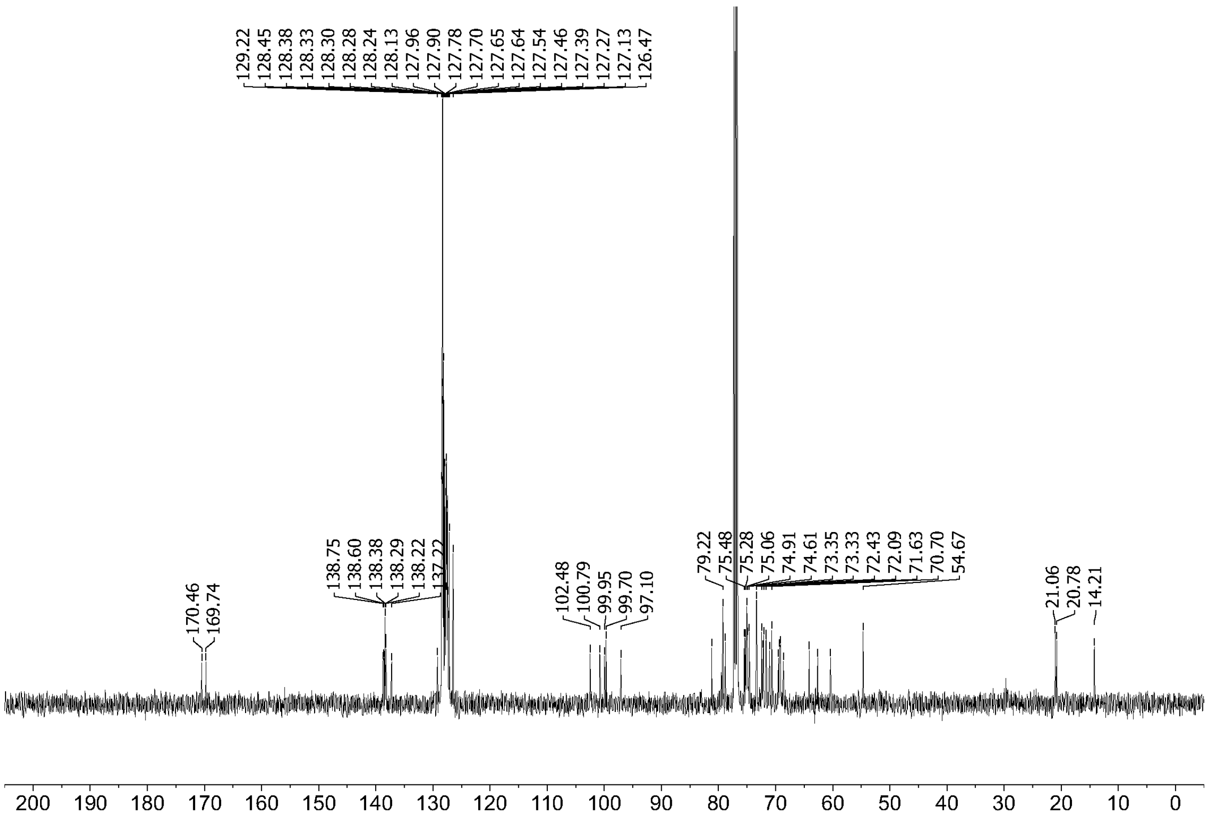
129.22
128.45
128.38
128.33
128.30
128.28
128.24
128.13
127.96
127.90
127.78
127.70
127.65
127.64
127.54
127.46
127.39
127.27
127.13
126.47
170.46
169.74
138.75
138.60
138.38
138.29
138.22
137.22
102.48
100.79
99.95
99.70
97.10
79.22
75.48
75.28
75.06
74.91
74.61
73.35
73.33
72.43
72.09
71.63
70.70
54.67
21.06
20.78
14.21
200 190 180 170 160 150 140 130 120 110 100 90 80 70 60 50 40 30 20 10 0

**Methyl α-D-glucopyranosyl-(1,3)-α-D-mannopyranosyl-(1,2)-α-D-mannopyranosyl-(1,2)-α-D-mannopyranoside (7)**

$^{1}$H NMR (500 MHz, $D_2O$)

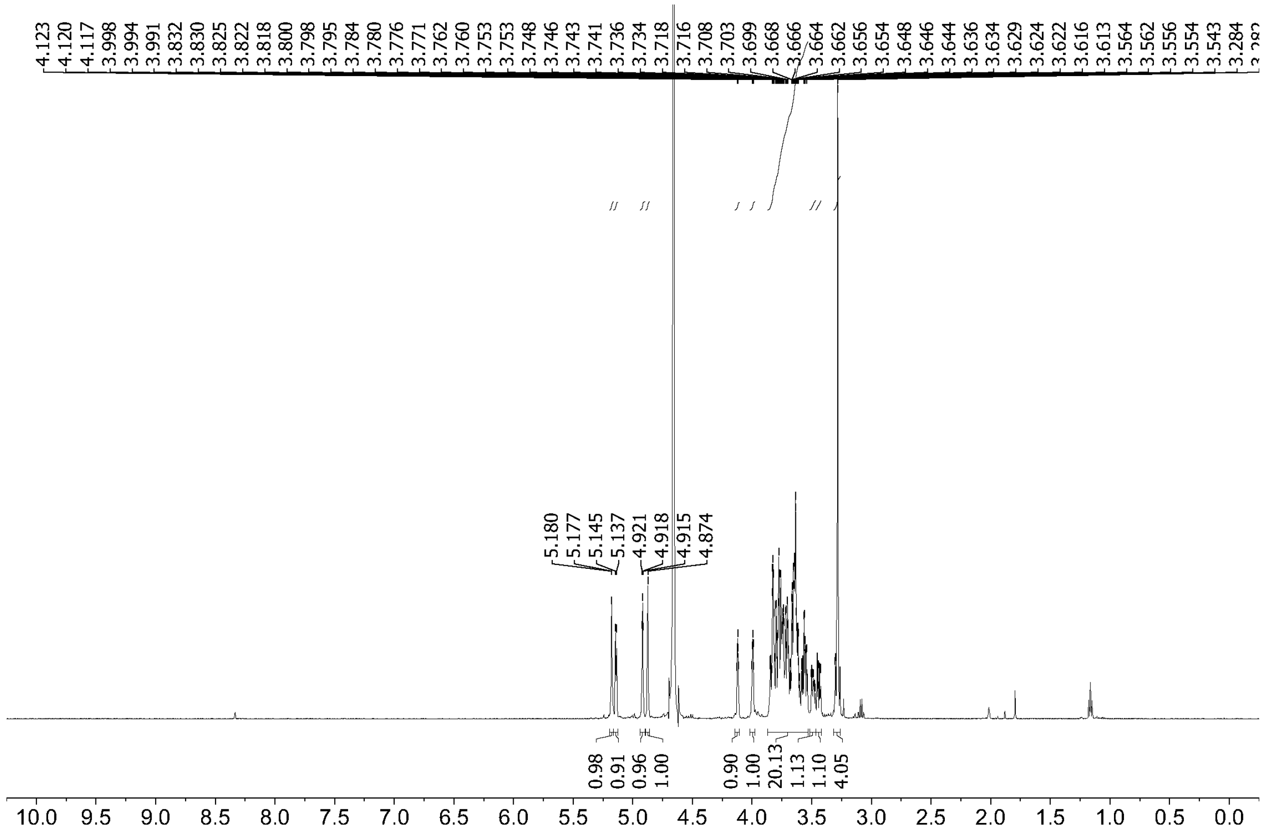

$^{13}$C NMR (125 MHz, $D_2O$)

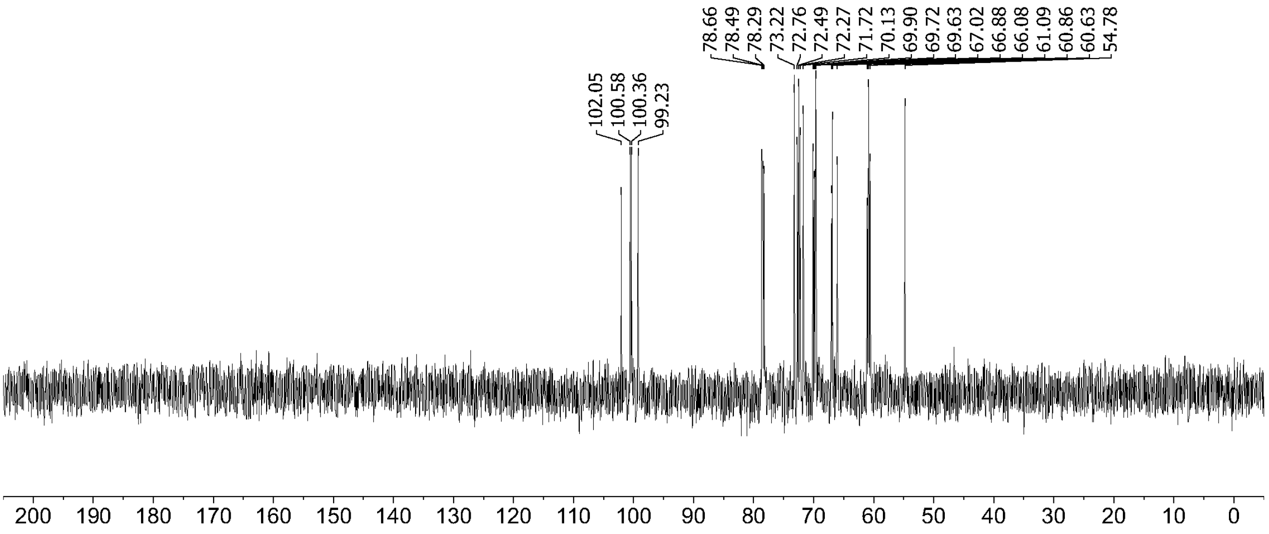

$^{1}H$-$^{1}H$ COSY

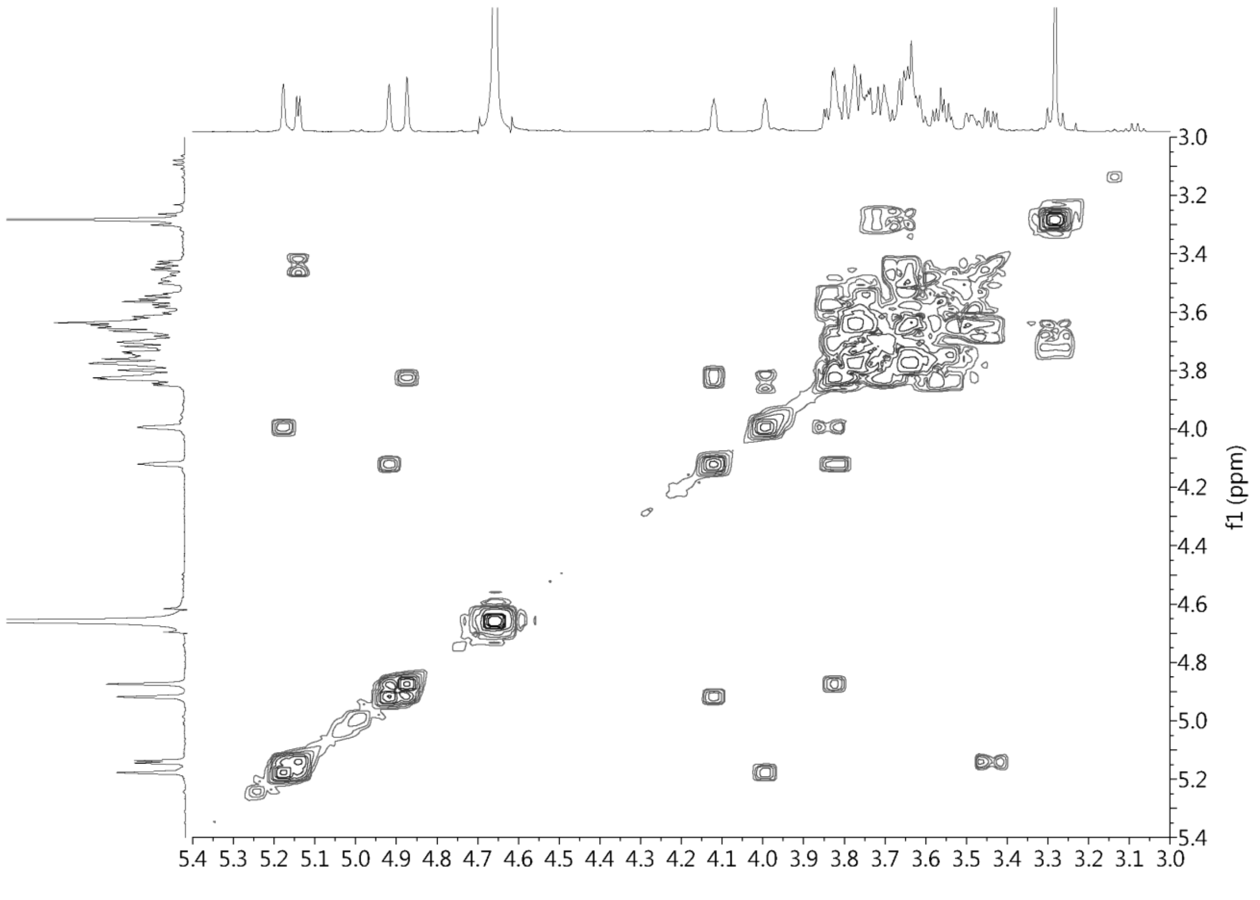

$^{1}H$-$^{13}C$ COSY (HSQC)

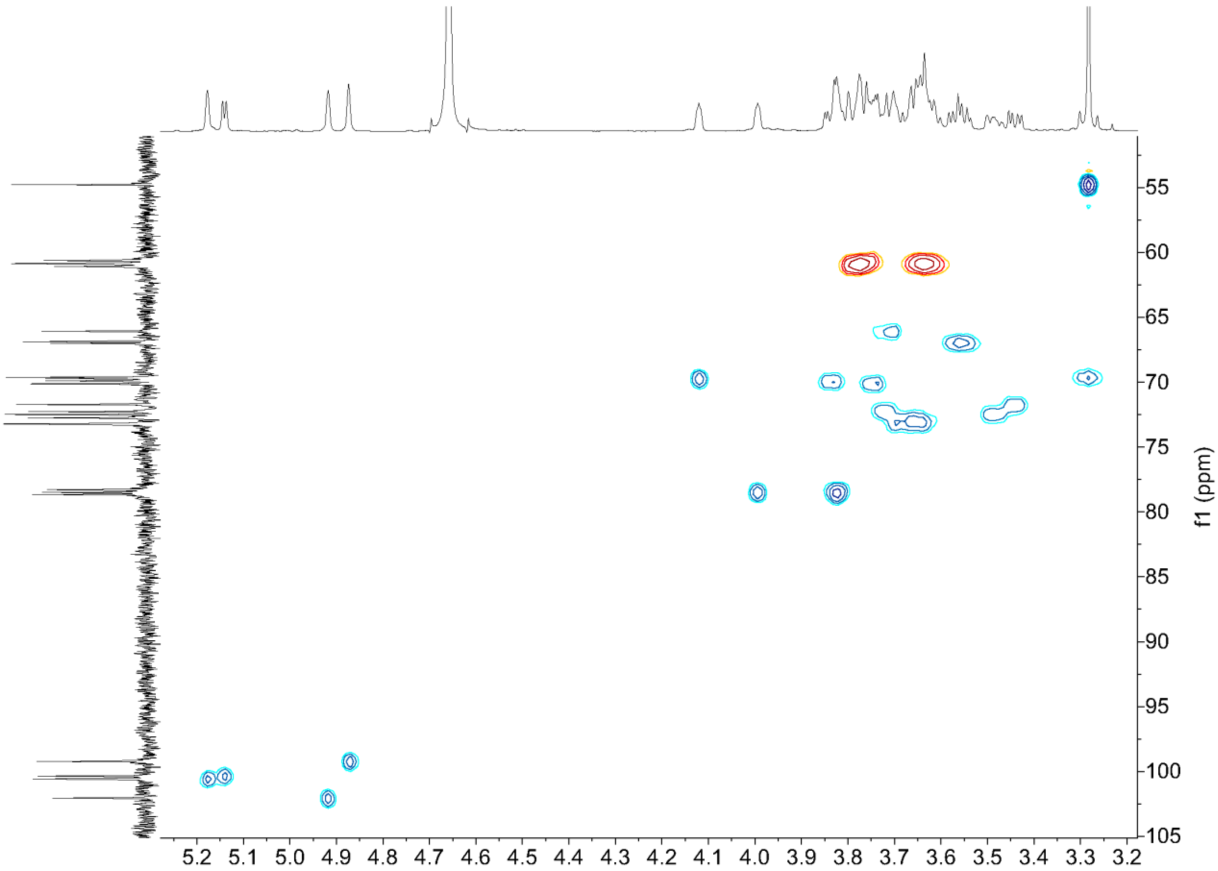

f1 (ppm)
55
60
65
70
75
80
85
90
95
100
105
5.2 5.1 5.0 4.9 4.8 4.7 4.6 4.5 4.4 4.3 4.2 4.1 4.0 3.9 3.8 3.7 3.6 3.5 3.4 3.3 3.2

**3-*O*-Cyclohexylmethyl-1,2:5,6-di-*O*-isopropylidene-α-D-glucofuranose (8)**

$^{1}$H NMR (500 MHz, $CDCl_3$)

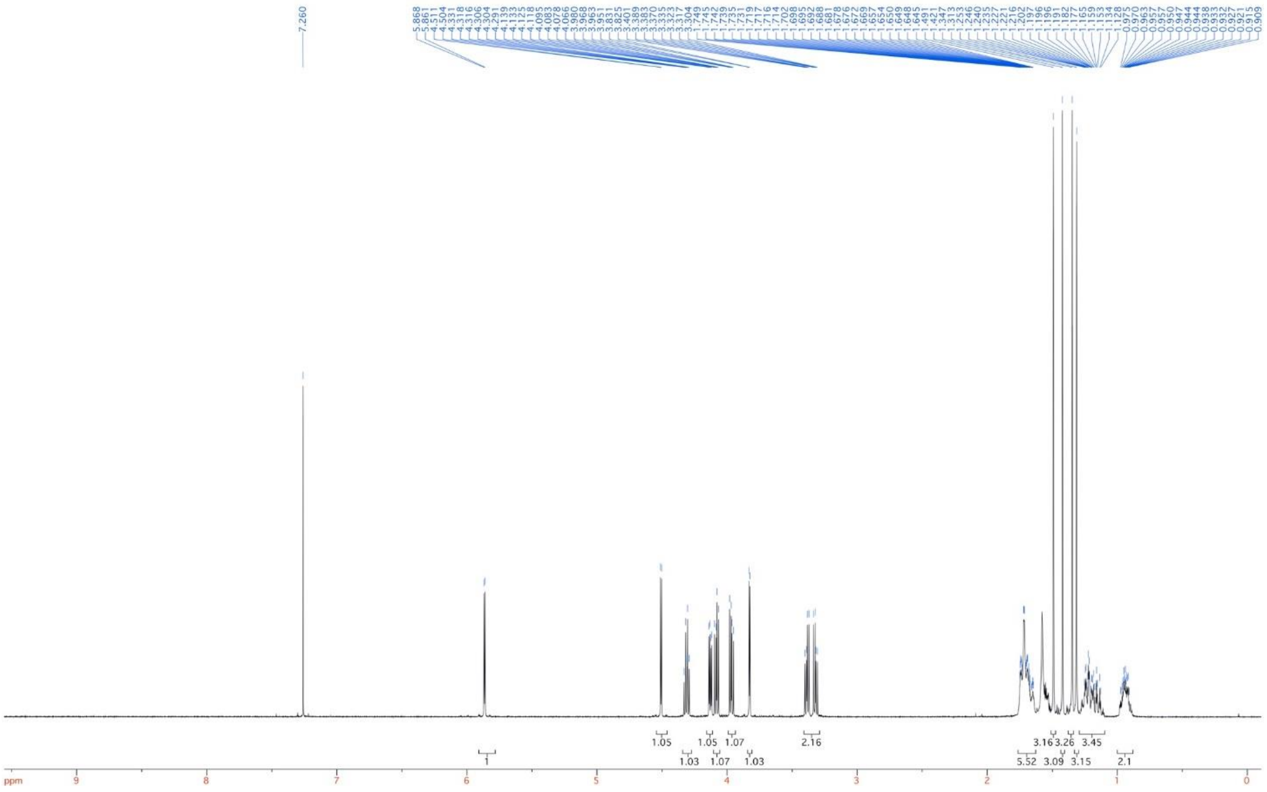

$^{13}$C NMR (125 MHz, $CDCl_3$)

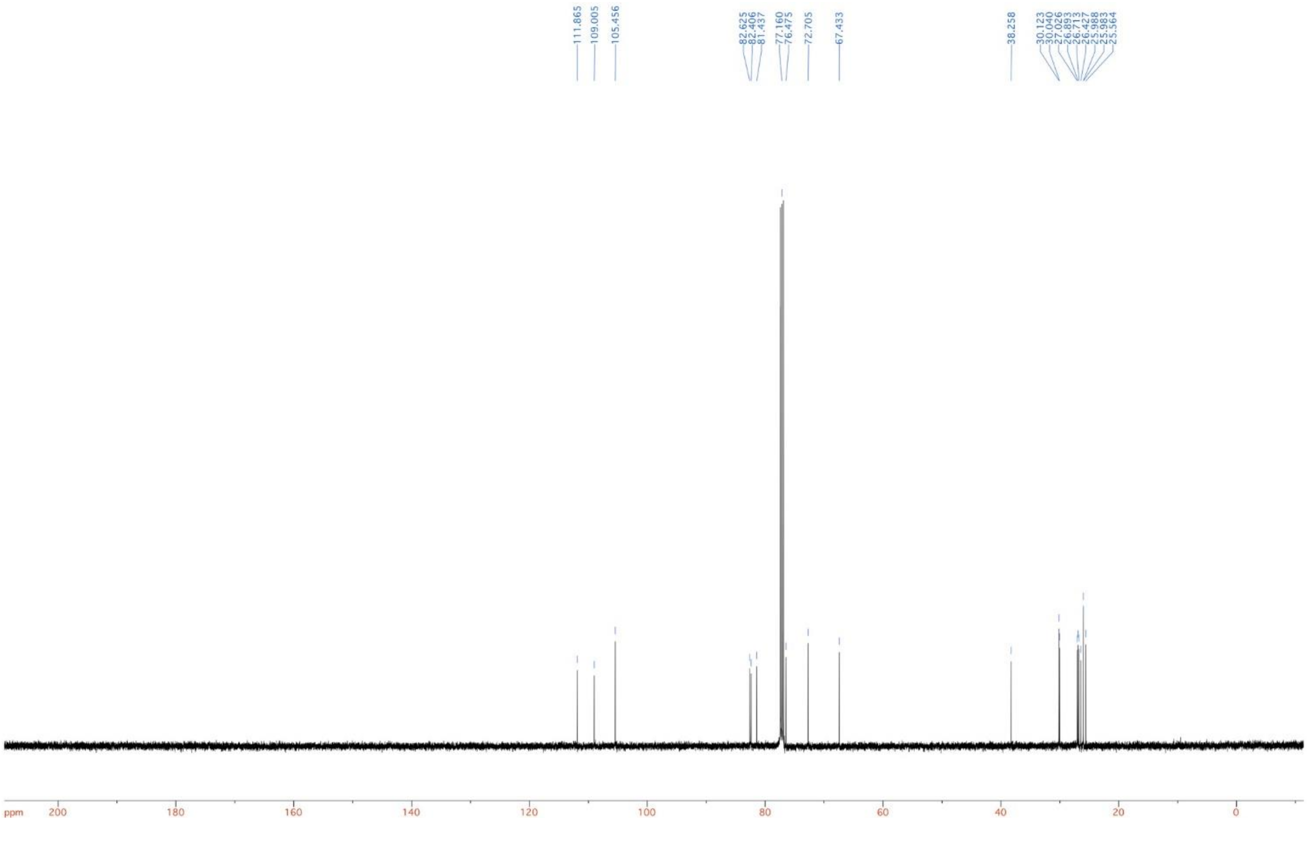

**Methyl 3-*O*-cyclohexylmethyl-α/β-D-glucopyranoside (10)**

$^{1}$H NMR (500 MHz, $d_4$-MeOH):

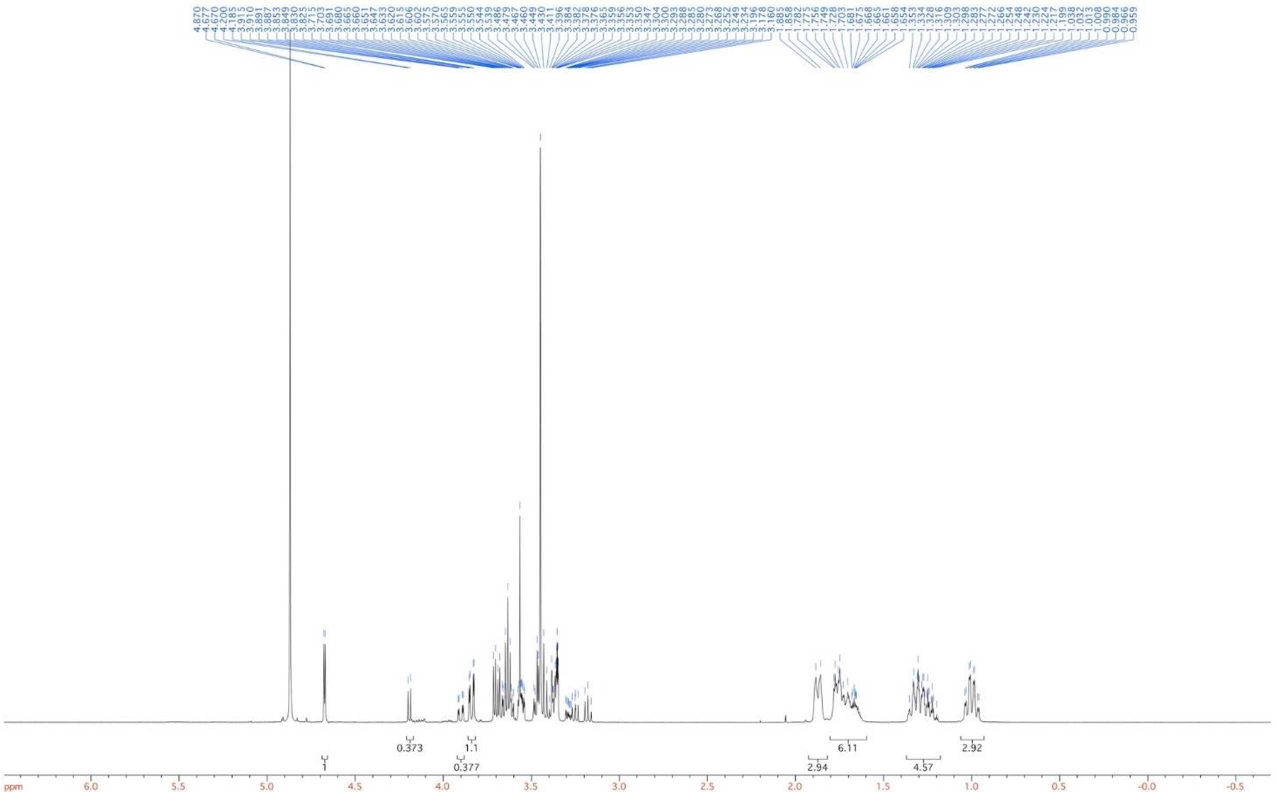

$^{13}$C NMR (125 MHz, $d_4$-MeOH)

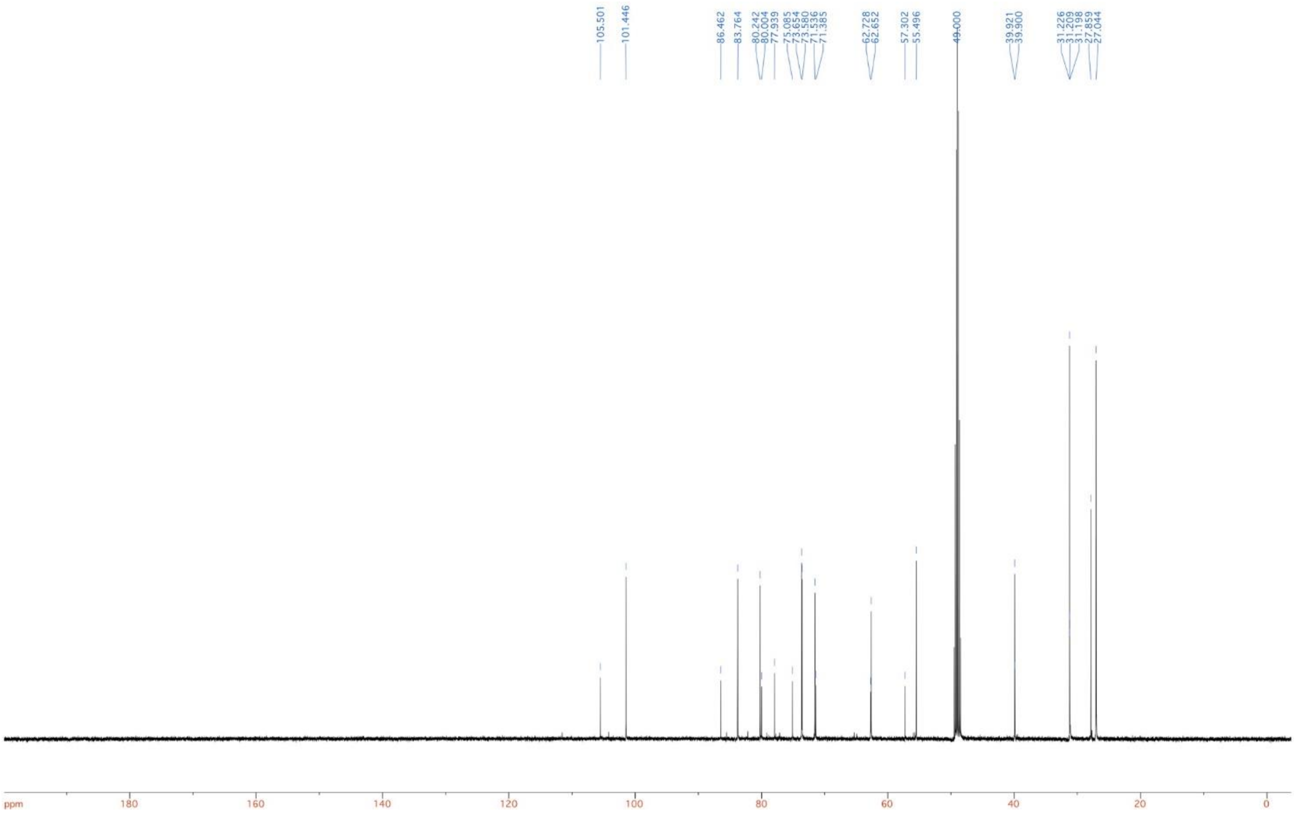
105.501
101.446
86.462
83.764
80.242
80.004
77.939
75.085
73.654
73.580
71.536
71.385
62.728
62.652
57.302
55.496
49.000
39.921
39.900
31.226
31.209
31.198
27.859
27.044
ppm
180
160
140
120
100
80
60
40
20
0

**Methyl 2,4,6-tri-*O*-benzyl-3-*O*-cyclohexylmethyl-α/β-D-glucopyranoside (11)**

$^{1}$H NMR (500 MHz, $CDCl_3$):

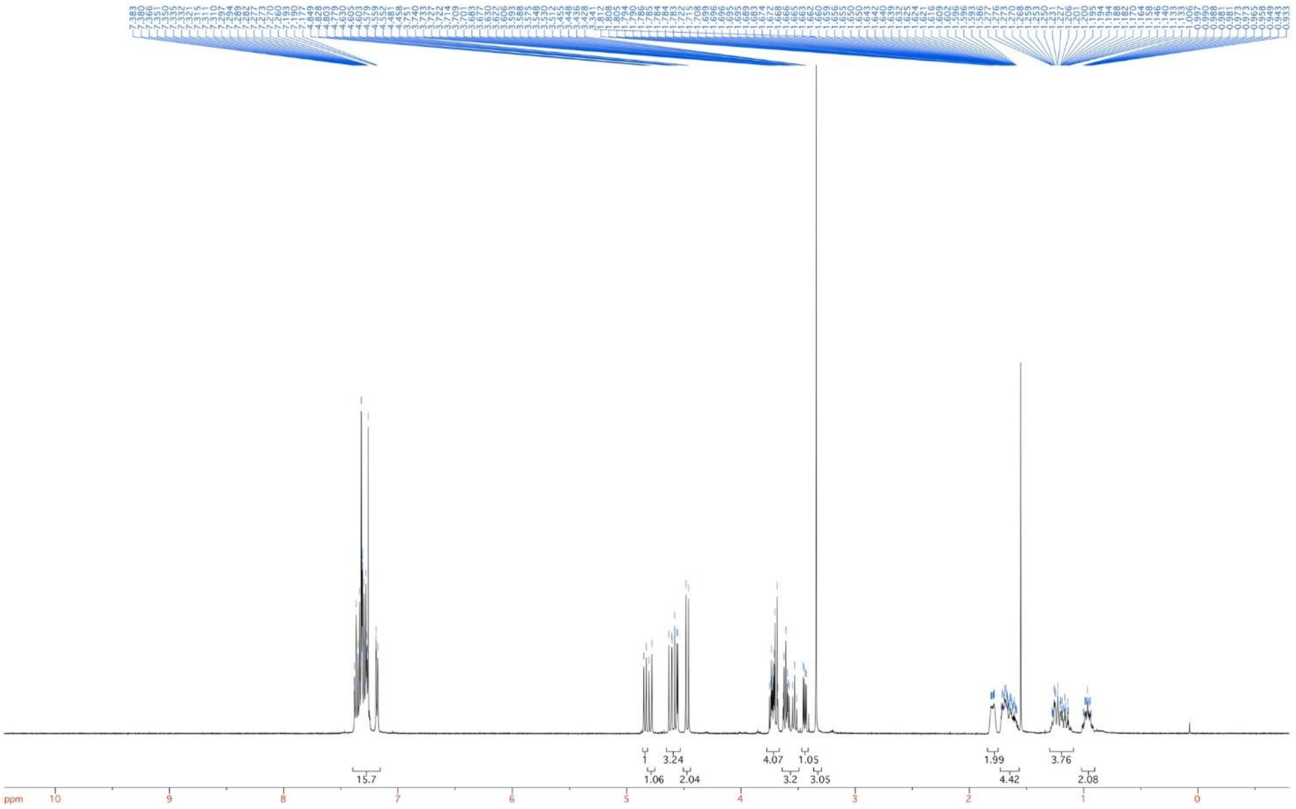

$^{13}$C NMR (125 MHz, $CDCl_3$)

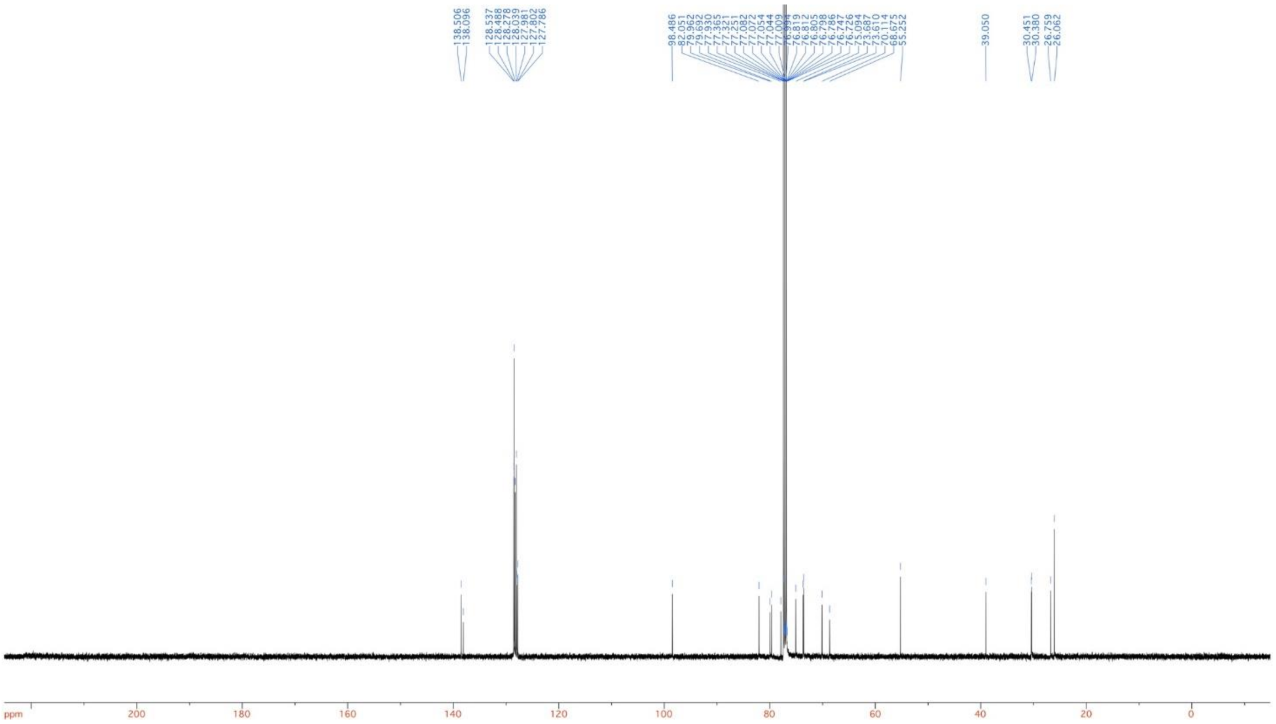

**1,6-Di-*O*-acetyl-2,4-di-*O*-benzyl-3-*O*-cyclohexylmethyl-α/β-D-glucopyranoside (12)**

$^{1}$H NMR (500 MHz, $CDCl_3$)

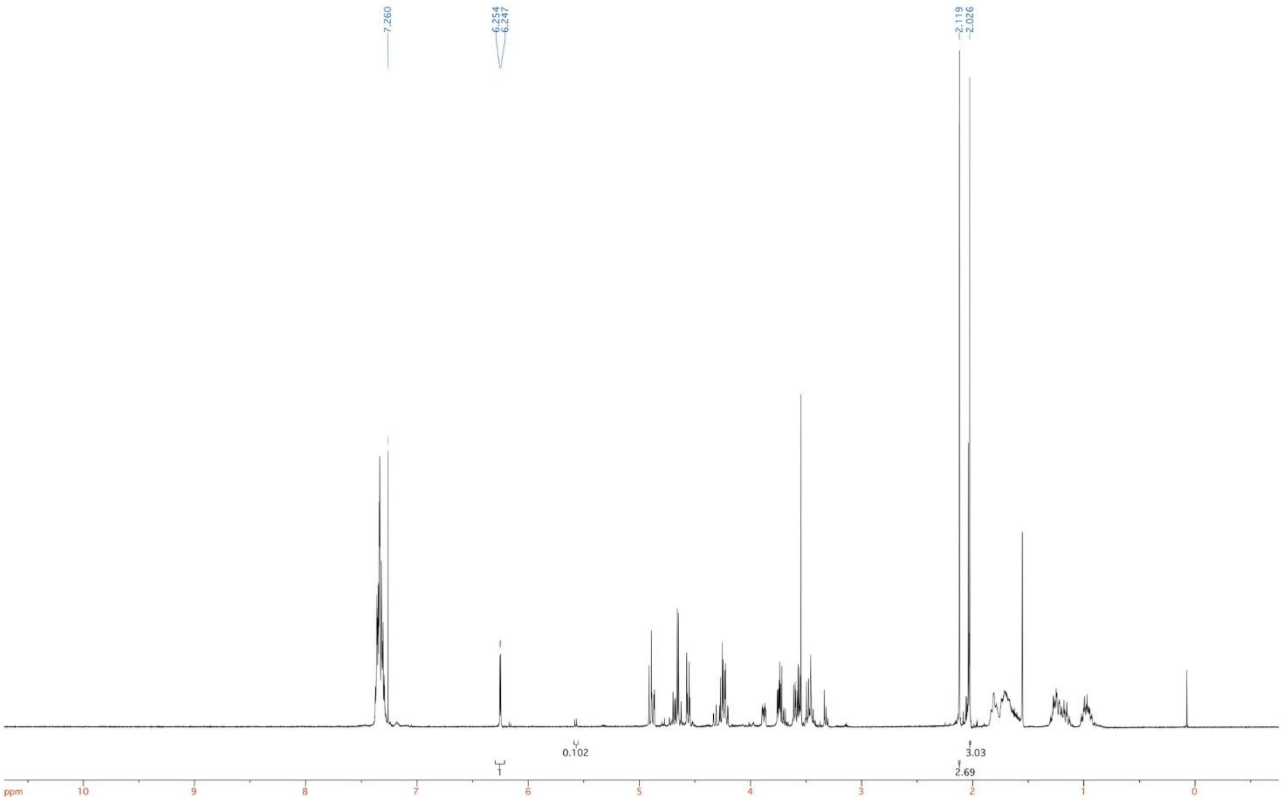

$^{13}$C NMR (125 MHz, $CDCl_3$)

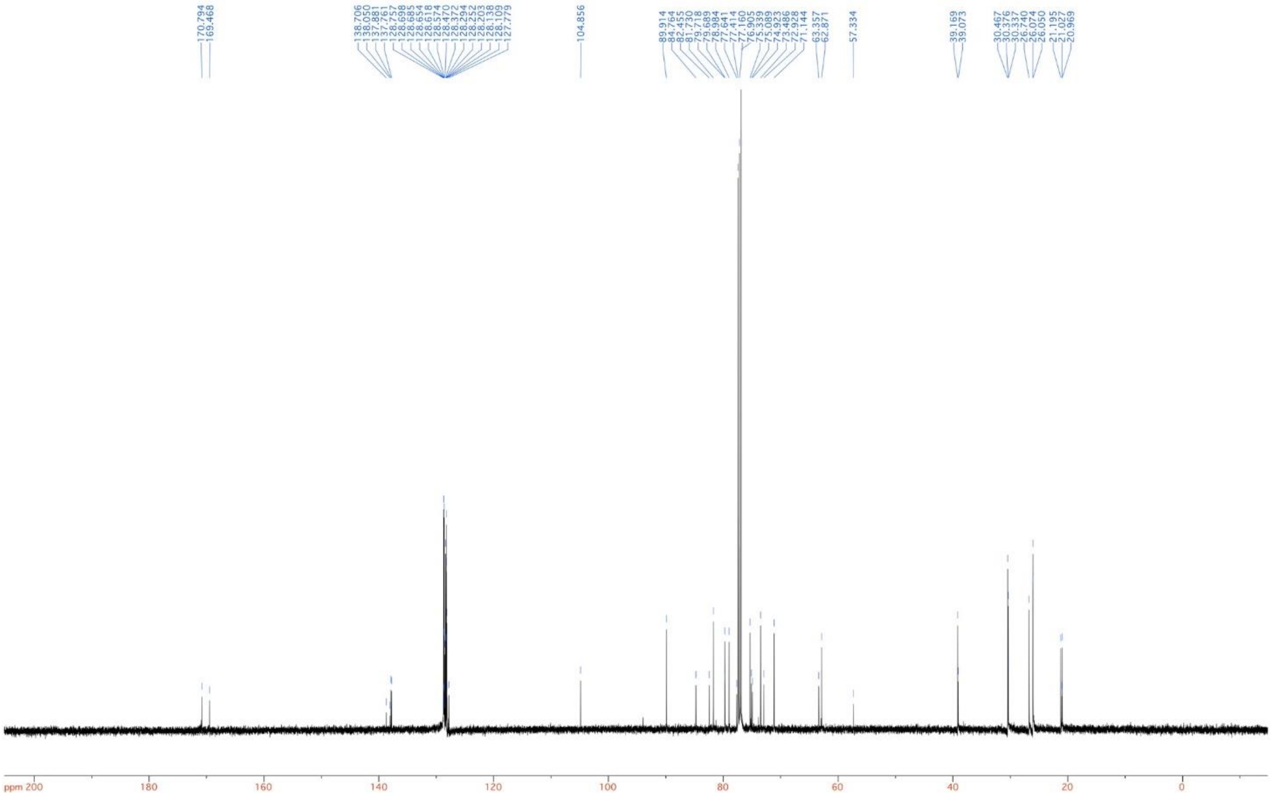

**3-*O*-(6’-*O*-Acetyl-2’,4’-di-*O*-benzyl-3’-*O*-cyclohexylmethyl-α-D-glucopyranosyl)-(4,6-*O*-benzylidene-*N*-benzyloxycarbonyl-isofagomine (15)**

$^{1}H$ NMR (500 MHz, $CDCl_3$):

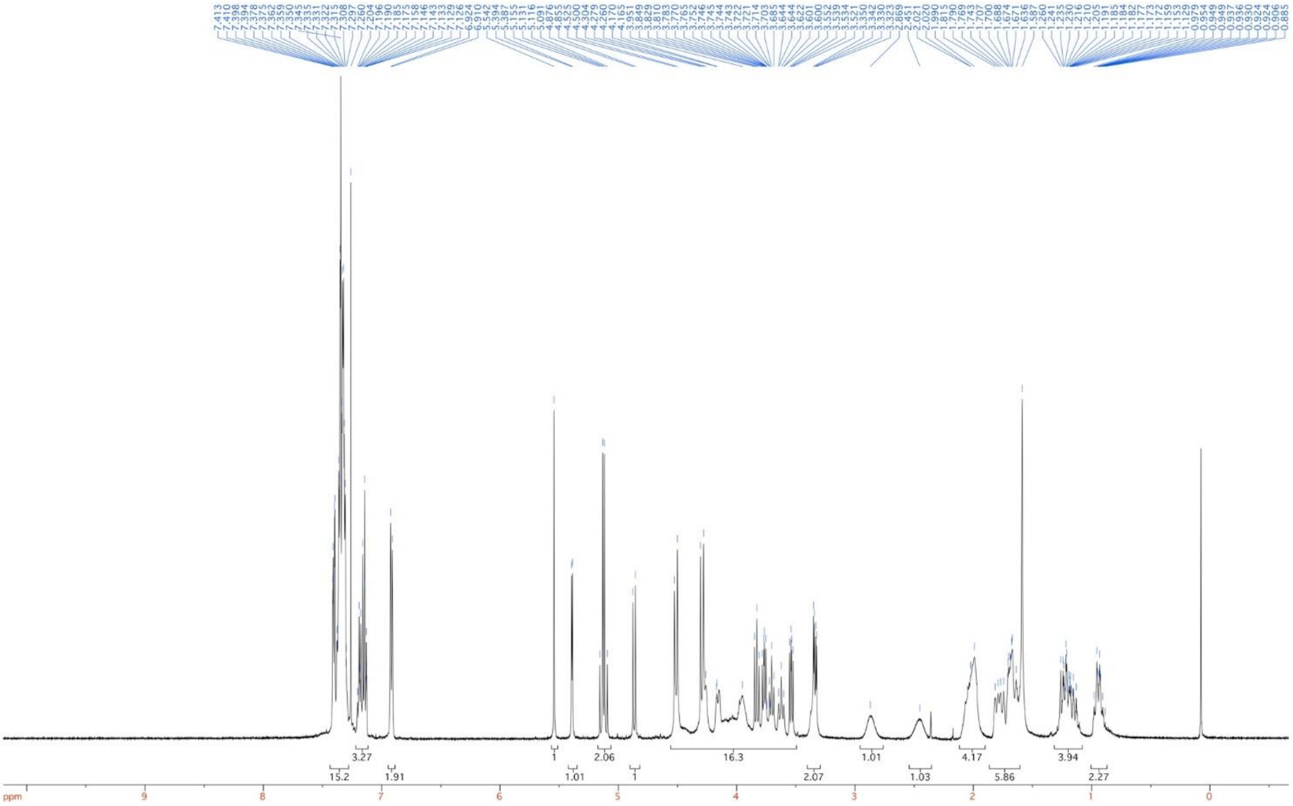

$^{13}C$ NMR (125 MHz, $CDCl_3$)

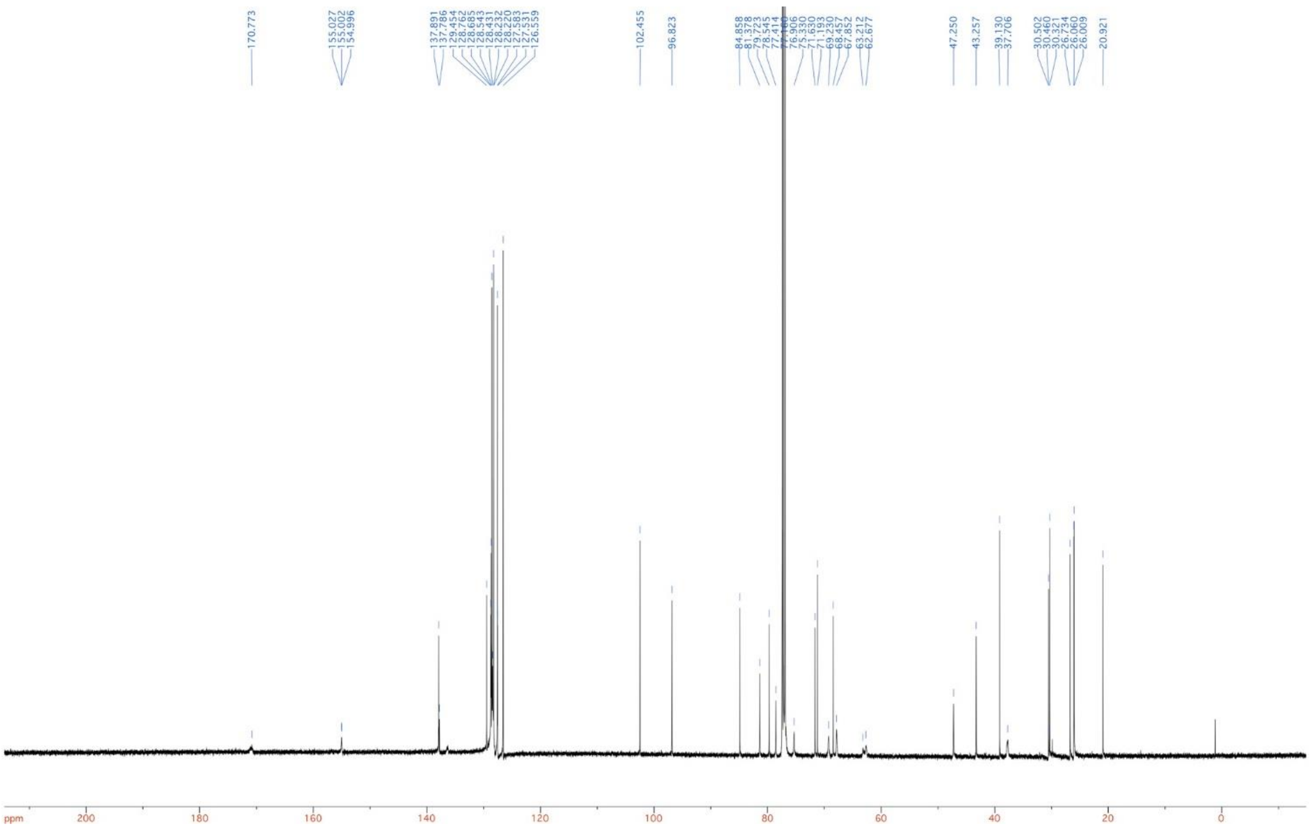

**(3*R*,4*R*,5*R*)-3-(3'-*O*-Cyclohexylmethyl-α-D-glucopyranosyloxy)-4-hydroxy-5-(hydroxymethyl)-piperidine (16, CyMe-GlcIFG)**

$^{1}$H NMR (500 MHz, $D_2O$/$d_4$-MeOH 1:1)

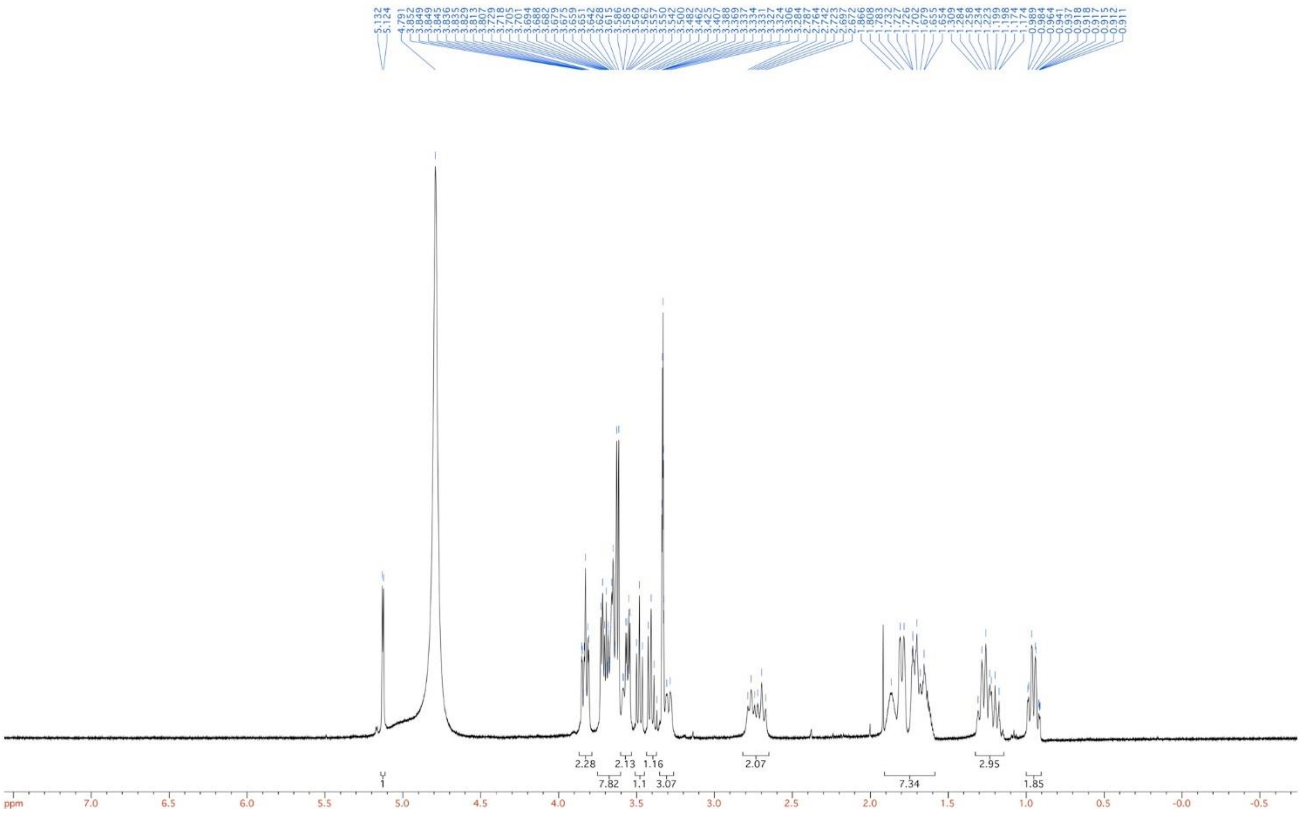

$^{13}$C NMR (125 MHz, $D_2O$/$d_4$-MeOH 1:1)

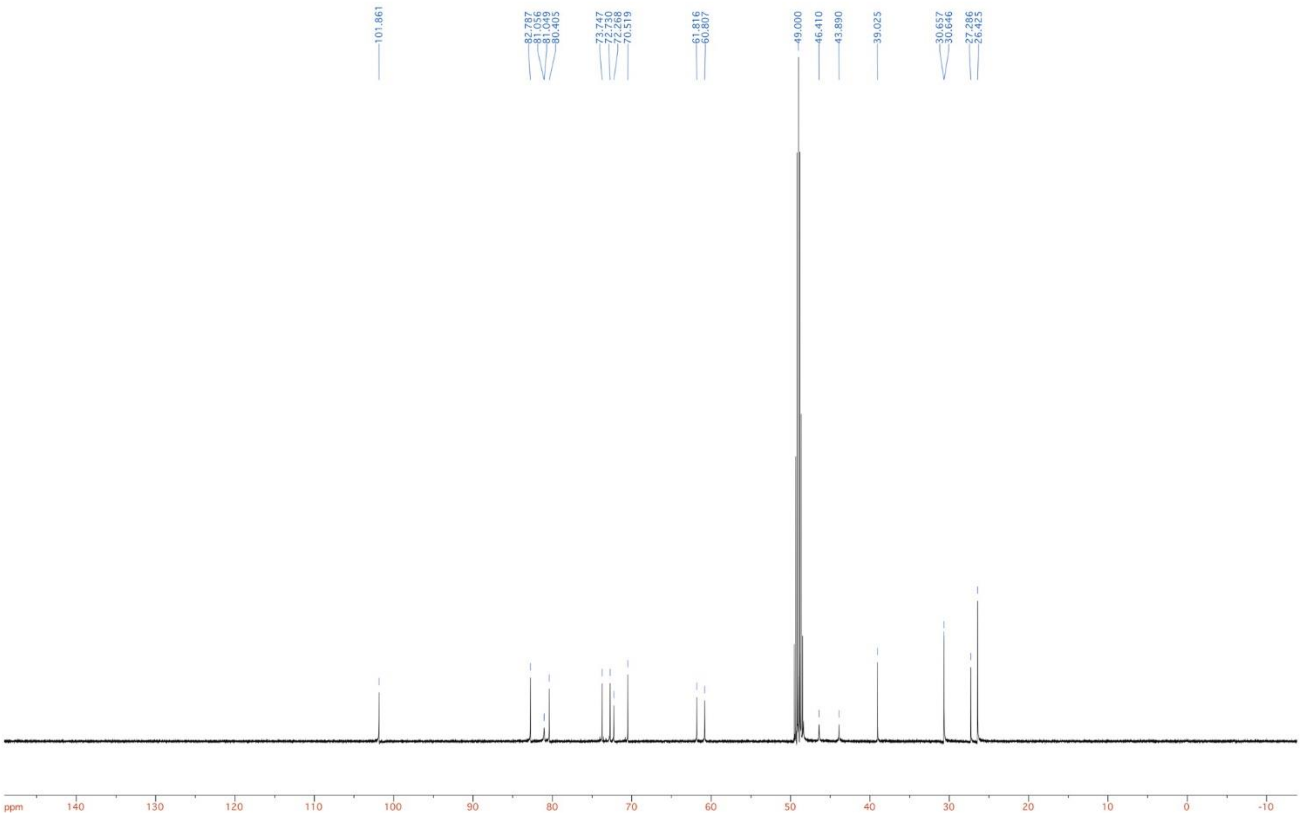
101.861
82.787
81.056
81.049
80.405
73.747
72.730
72.268
70.519
61.816
60.807
49.000
46.410
43.890
39.025
30.657
30.646
27.286
26.425
ppm
140
130
120
110
100
90
80
70
60
50
40
30
20
10
0
-10
